# Comparative Evaluation of Assumption Lean Community Detection Methods for Human Connectome Networks

**DOI:** 10.1101/2025.11.13.688333

**Authors:** Ayoushman Bhattacharya, Nilanjan Chakraborty, Jiaxin Cindy Tu, Xintian Wang, Donna Dierker, Andy Eck, Jed T. Elison, Soumen Lahiri, Adam Eggebrecht, Muriah D. Wheelock

## Abstract

Community detection on resting-state functional connectivity provides a principled lens on mesoscale organization in functional brain networks. The choice of community count *K*, lacks a standardized or principled guideline. We conducted a benchmark of three approaches that do not a priori restrict communities to an assortative mesoscale topology: the Weighted Stochastic Block Model, Spectral Clustering, and K-means Clustering. Performance was assessed on synthetic networks with ground truth and on three neuroimaging cohorts which included adult and infant datasets. We compared several strategies for selecting *K*, including the silhouette index and other approaches commonly used in the literature. These were evaluated alongside a likelihood-based criterion for the weighted stochastic block model that employs bootstrap confidence intervals for differences in log-likelihood between successive values of *K*. For the synthetic networks, WSBM and Spectral Clustering correctly identified the true number of communities, whereas K-means Clustering did not. In adult datasets, most indices did not yield a unique optimum, whereas the likelihood-based criterion selected *K* = 11, which is consistent with established sensory and association systems. In infants and toddlers, the same procedure supports a larger *K* around 15 and reveals developmentally distinct mesoscale architecture, including anterior and posterior subdivisions within default mode and frontoparietal systems.

## Introduction

Large-scale functional systems are detectable early in life, and aspects of an adult-like network scaffold are already present during infancy. In particular, primary sensory and motor networks exhibit relatively mature topological organization early in development, and selective long-range functional connections can be observed even in neonates (Derman et al., 2025; Gao, Alcauter, Elton, et al., 2015; Gao, Alcauter, Smith, et al., 2015; Grayson & Fair, 2017; Moore et al., 2024; Sylvester et al., 2023). These findings have supported the view that the broad functional architecture of the infant brain resembles that of the adult brain. In contrast, age-specific data-driven analyses indicate that adult-derived parcellations fit neonatal and infant connectivity incompletely, that higher-order association systems remain spatially fragmented, and that infant-specific solutions better capture early-life functional organization (Eggebrecht et al., 2017; Eyre et al., 2021; Kardan et al., 2022; Myers et al., 2024; Tu et al., 2026; Tu, Myers, et al., 2025; Tu, Wang, et al., 2025; Wang et al., 2023) . Thus, the presence of an early adult-like scaffold does not necessarily imply that the mesoscale functional organization of the developing brain is equivalent to that of the adult brain. An important unresolved question is whether early-life functional organization is adequately represented by an adult-derived community structure or instead exhibits a developmentally distinct mesoscale partition, including differences in the number, granularity, and arrangement of functional communities. Addressing this question is especially challenging because task-based validation of functional networks is limited during early development.

Testing whether early-life mesoscale organization differs from adult organization requires community-detection methods that do not prespecify an adult network atlas and that are sufficiently flexible to recover more than strictly assortative structure. Clustering algorithms widely used in the neuroimaging literature often make assumptions about the organization of brain networks. While there is a broad consensus among neuroscientists that the human connectome is predominantly assortative, some research has suggested that many but not all communities are assortative or may have different structures, namely core and periphery (Betzel, Bertolero, et al., 2018; Faskowitz & Sporns, 2020). The most commonly used community detection algorithms in the neuroimaging literature, namely modularity maximization (Blondel et al., 2008; Newman & Girvan, 2004), Infomap (Eggebrecht et al., 2017; Gordon et al., 2016; Kardan et al., 2022; Power et al., 2012; Rosvall & Bergstrom, 2008; Tu, Myers, et al., 2025; Wheelock et al., 2019), von Mises-Fisher distribution (Yeo, et al., 2011), and gradient-weighted Markov Random Field model (Schaefer et al., 2018), works on the assumption that sub-networks are internally dense and externally sparse. This may place a constraint on the connectome’s meso-scale structure, favoring a more assortative organization (Betzel, Bertolero, et al., 2018; Betzel, Medaglia, et al., 2018; Faskowitz & Sporns, 2020). Alternatively, others (Faskowitz et al., 2018; Faskowitz & Sporns, 2020; Tooley et al., 2022) suggested that Weighted Stochastic Block Model (WSBM), allows a greater flexibility to detect a diverse range of network structures, including modular, disassortative (Betzel, Bertolero, et al., 2018), and core-periphery (Noori et al., 2017; Pavlovic et al., 2014) configurations. Brooks and colleagues further demonstrated that the performance and agreement of community-detection methods depend on network topology, underscoring the importance of evaluating findings across multiple analytical approaches (Brooks et al., 2024).

Determining the optimal number of communities in the human connectome remains an open challenge for several reasons. For widely used approaches such as modularity maximization, a single modular partition may represent only one of several plausible solutions, and the resulting partitions may also be affected by the resolution limit (Fortunato & Barthélemy, 2007; Fortunato & Hric, 2016; Peel et al., 2017). Conversely, other approaches, such as the asymptotical surprise algorithm, can identify extremely small communities that may not be biologically meaningful (Nicolini et al., 2017). Previous studies have also used post hoc criteria—including Normalized Mutual Information-based stability analysis (Yeo et al., 2011), the zRand index (Akiki & Abdallah, 2019; Doron et al., 2012; Traud et al., 2011), Variation of Information distance (Faskowitz & Sporns, 2020; Meilă, 2007), and the silhouette index—to determine the optimal number of communities. However, these criteria can produce different preferred community numbers or assignments depending on the clustering algorithm and analytical setting (Sanchez-Rodriguez et al., 2021; Yeo et al., 2011). This uncertainty is particularly consequential in developmental connectomes: reported infant network solutions range from approximately 6 to 23 communities (Sylvester et al., 2023; Tu et al., 2026), although these values are not directly comparable because they variously reflect prespecified templates, retained ICA components, density-dependent community solutions, stability-guided choices, and deliberately coarse or fine representations. Consequently, it remains unclear whether differences among reported infant network organizations reflect neurodevelopmental biology, analytical choices, or both. Imposing an adult-derived number of communities may also obscure subdivisions or organizational features specific to early development. Complementing Brooks and colleagues, who examined topology-dependent differences among community-detection methods, we therefore focus on the principled selection of (*K*) and its implications for mesoscale organization across adult and early-life connectomes (Brooks et al., 2024).

In this article, our primary objective is to test the hypothesis that the mesoscale functional organization of the early-life brain is developmentally distinct from that of the adult brain, despite the presence of an early adult-like functional scaffold. We investigate this hypothesis by asking whether a common, data-driven community-detection framework identifies differences in the preferred number, granularity, and arrangement of functional communities between adult and early-life connectomes, and whether these developmental features converge across multiple analytical approaches. To this end, we evaluate three community-detection algorithms that do not a priori constrain the inferred communities to an assortative mesoscale topology: the weighted stochastic block model (WSBM) (Aicher et al., 2013, 2015), spectral clustering (Abbe, 2018; Cribben & Yu, 2017; Rohe et al., 2011), and K-means clustering (Thirion et al., 2014; Tononi et al., 1998). We first use synthetic networks with known ground-truth structure to evaluate alternative strategies for selecting the number of communities. We then apply these methods to adult and early-life functional-connectivity data, assess the reproducibility and consensus of the inferred partitions, and compare them with established adult and infant cortical atlases defined using the Infomap community-detection algorithm (Gordon et al., 2016; Kardan et al., 2022). For the early-life data, we additionally benchmark the inferred community organization using Louvain modularity maximization, providing a complementary comparison with a widely used assortativity-based method. Through this convergent analytical framework, we seek to determine whether the developing connectome is best characterized as an adult-like organization or as a related but developmentally distinct mesoscale architecture.

## Methods

### Synthetic human connectome

To assess the performance of the clustering algorithms in the situation where the true number of communities are known, synthetic brain functional connectivity with a pre-specified network count and size were generated for testing the ability of different clustering models to identify the correct number of communities. Using a known probability distribution and a set of parameters based on the Human Connectome Project (HCP) and Baby Connectome Project (BCP) functional connectivity (FC) data (details described under in vivo datasets), we generated the synthetic brain networks. Specifically, we used WSBM to create a functional connectivity matrix for various choices of community and number of ROI as described in Table S1 in the Supplementary Materials. To mimic the variability of a human brain connectome, we used Infomap communities defined by the Gordon atlas (Gordon et al., 2016). The mean and variance of FC within and between the communities were chosen based on the distribution observed in HCP and BCP data (Fig. S1), and FC between pairs of ROIs were generated from a normal distribution (Fig. 1). For each ground-truth value *K* = 5, *K* = 7, and *K* = 10, we generated 100 independent FC matrices under each of the HCP- and BCP-informed mean and variance settings. For each FC matrix and candidate number of communities, each clustering algorithm was run using 1,000 initializations, yielding 100,000 runs per algorithm per candidate value within each simulation setting. All valid WSBM solutions were retained. For each criterion, we then recorded the percentage of the 100 matrices for which each candidate value of *K* was selected; these selection frequencies are displayed as heatmaps (Figs. 3.e and 3.f).

**Figure 1.**
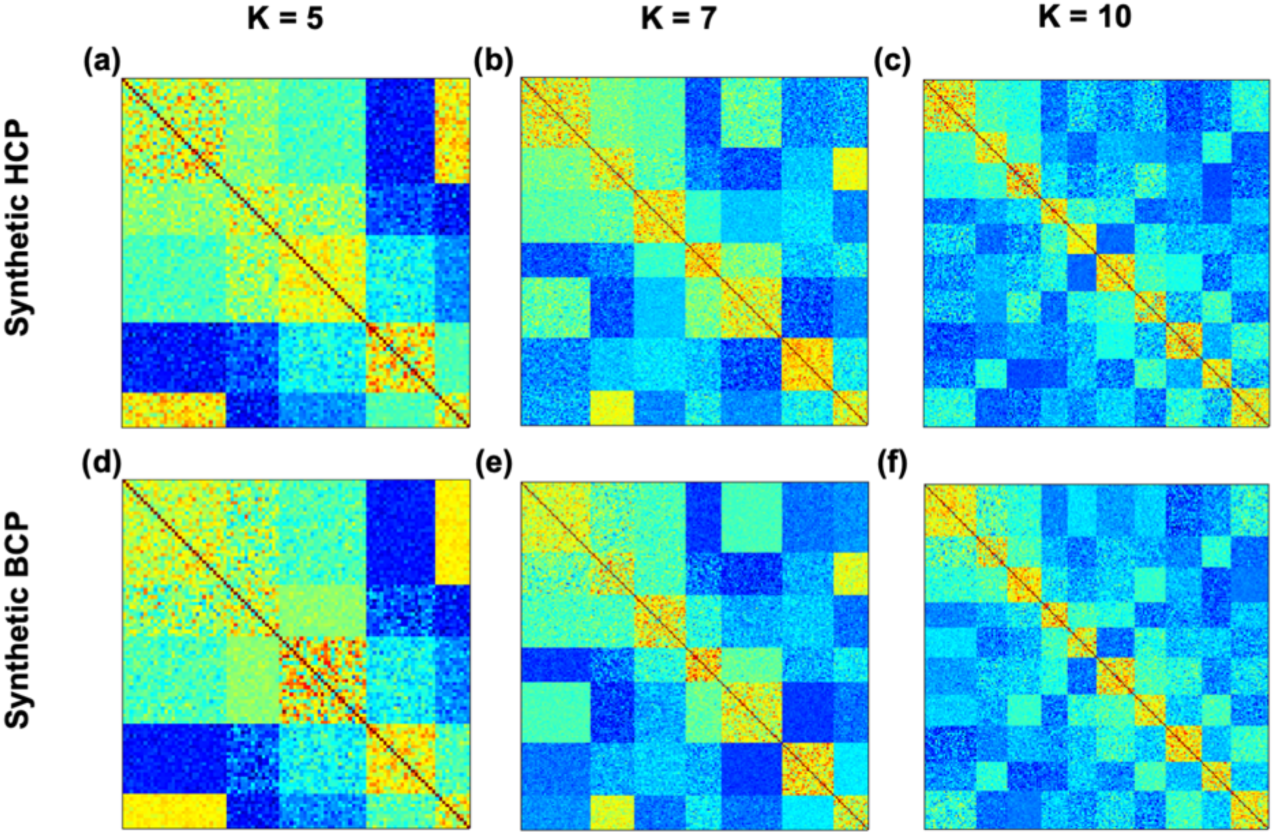
Illustration of a randomly selected simulated functional connectivity (FC) matrix from 100 independent simulations for each setting. **(a,b,c)** Synthetic FC matrices generated using pairwise connectivity means and variances estimated from HCP data for (*K* = 5,7,10) communities, respectively. **(d,e,f)** Corresponding synthetic FC matrices generated using BCP data.

**Figure 2.**
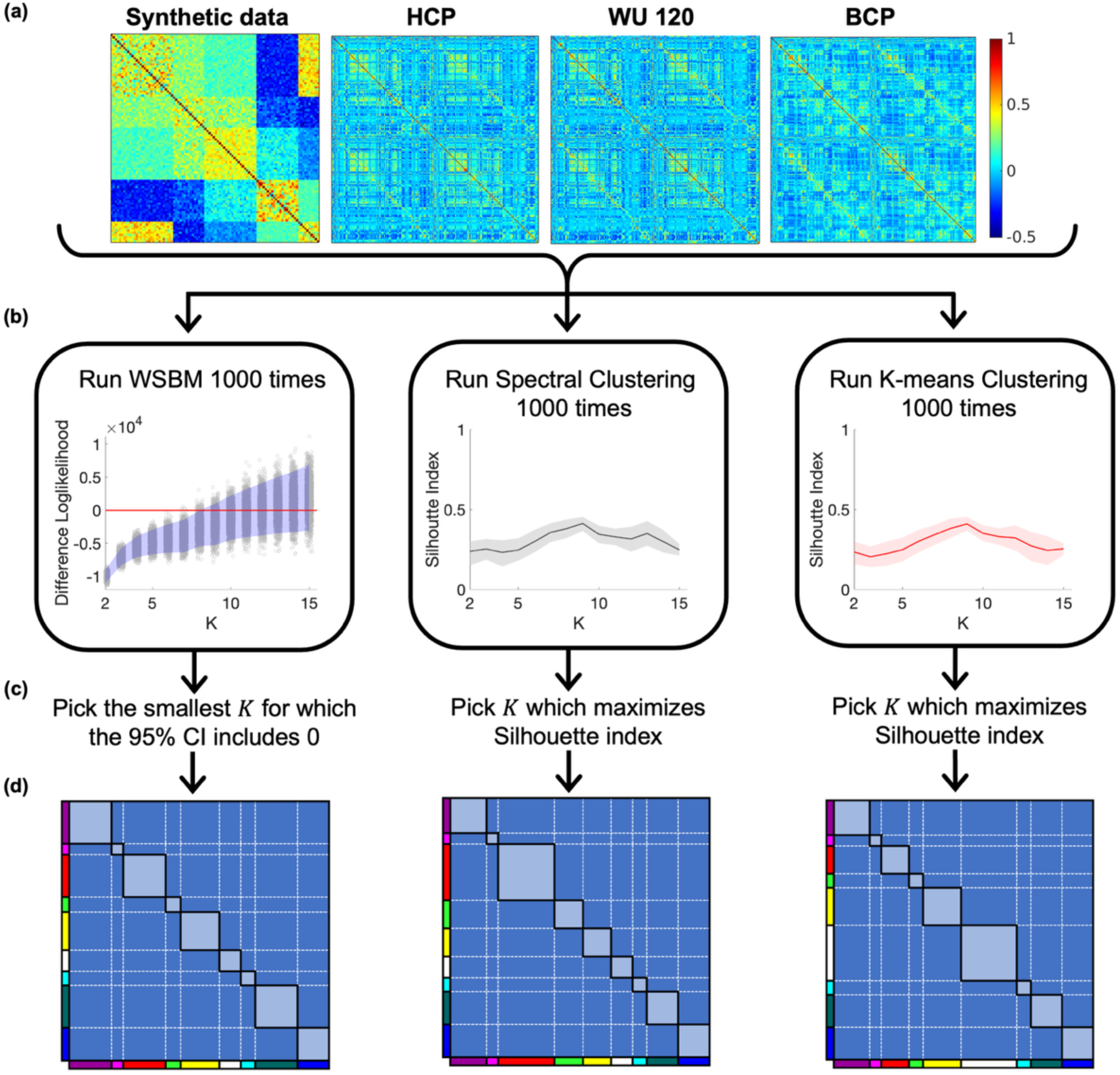
Diagrammatic representation of the methods and analyses for this study. **(a)** Average FC matrix of temporal correlation corresponding to the synthetic, HCP, WU 120 and BCP datasets. **(b)** For different choices of *K*, clustering is performed using all three clustering algorithms and an illustration of clustering performance are plotted. **(c)** Choice of optimal *K* for all three algorithms. **(d)** Illustration of final solutions based on optimal value of *K*. For WSBM, pick the solution corresponding to highest log-likelihood. For Spectral and K-means Clustering pick the solution corresponding to highest Silhouette index.

**Figure 3.**
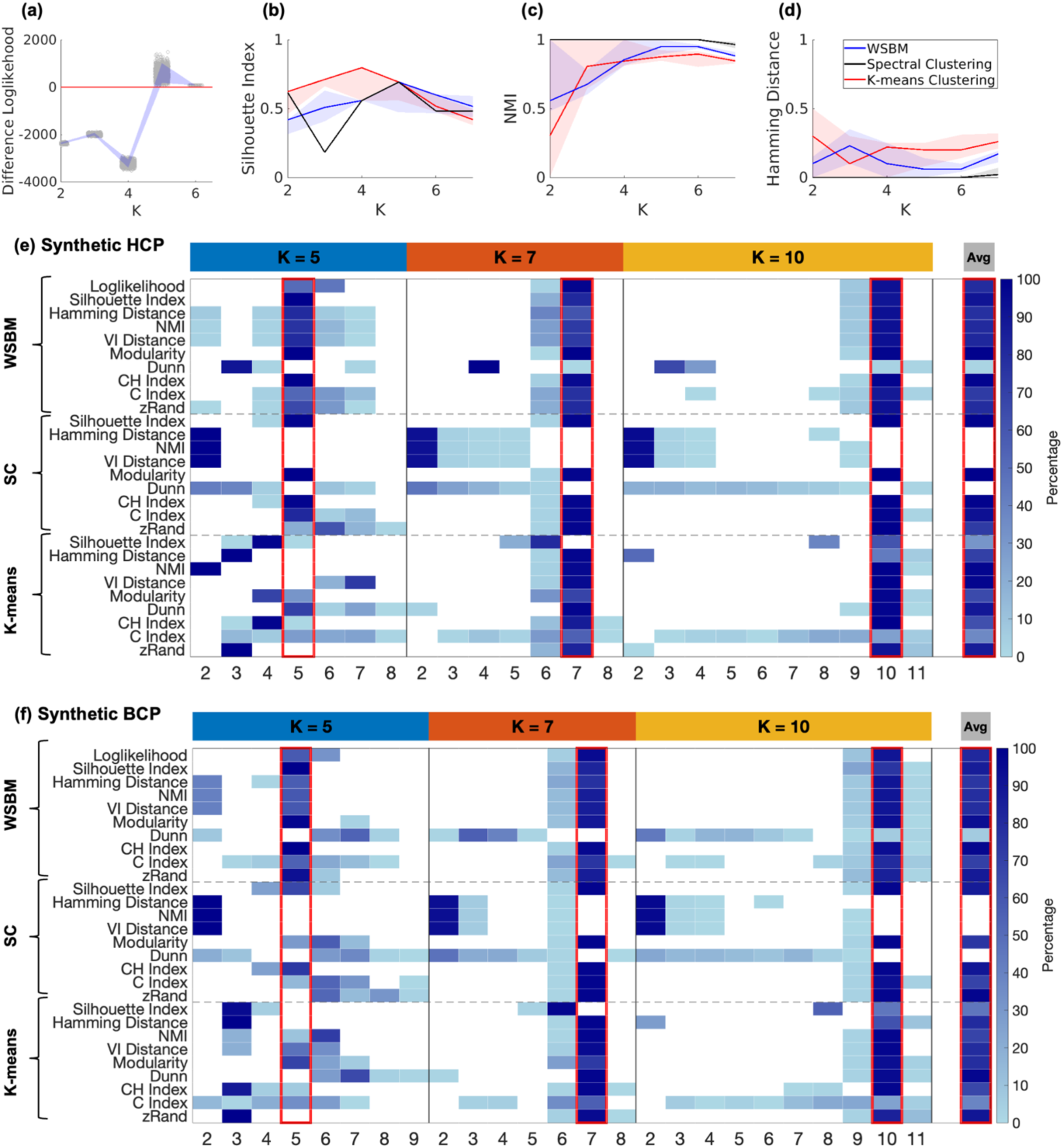
Synthetic data results. **(a–d)** Results from one randomly selected synthetic FC matrix among 100 independently generated matrices using HCP connectivity patterns with (*K* = 5) communities (Fig. 1a). **(a)** Illustration of bootstrap log-likelihood differences for WSBM, with the shaded region representing the 95% confidence interval. **(b,c,d)** Illustration of Silhouette index, NMI, and Hamming distance, respectively. The solid lines represent the median, and the shaded regions indicate the interquartile range across 1,000 clustering repetitions on the selected FC matrix. **(e,f)** Percentage of detected optimal numbers of communities for varying community sizes (Table S1), based on 100 independently generated synthetic FC matrices using connectivity patterns from HCP and BCP datasets, respectively (Fig. S1), with 1000 clustering repetitions performed on each FC matrix.

### Human Connectome Project dataset

We used the 3-Tesla (3T) resting-state fMRI data from the S1200 release of Human Connectome Project (HCP) dataset (Smith et al., 2013; Van Essen et al., 2012). The HCP study design involved recruiting twins and their family members to complete a series of MRI scans and behavioral assessments. We used 60-minute resting-state fMRI data collected across two days. Subjects were instructed to maintain a relaxed fixation on a white cross and not fall asleep (Smith et al., 2013). High-resolution T1-weighted images were obtained using a magnetization-prepared rapid acquisition gradient-echo (MP-RAGE) sequence, with a repetition time (TR) of 2.4 seconds and voxel dimensions of 0.7 × 0.7 × 0.7 mm. Additionally, BOLD contrast-sensitive images were collected using a gradient-echo echo-planar imaging (EPI) sequence, featuring a multiband factor of 8, a TR of 0.72 seconds, and voxel dimensions of 2 × 2 × 2 mm. These imaging procedures were conducted on each participant with a custom Siemens SKYRA 3.0T MRI scanner, which was equipped with a specialized 32-channel Head Matrix Coil designed for optimal signal acquisition. To enhance the robustness of the data, the HCP utilized imaging sequences with both left-to-right and right-to-left phase encoding directions. Each participant underwent one imaging session in each direction (Van Essen et al., 2012). The analysis included 965 healthy adults aged 22 to 35 with at least 10 minutes of low-motion data retained for scans after motion censoring at framewise displacement (FD) exceeding 0.04 mm. The data was minimally preprocessed (Glasser et al., 2013) and went through additionally custom processing involving nuisance regression, temporal filtering and frame censoring to further remove artifacts and mapped to cortical surfaces (32k-fsLR) as described in prior publications (Li et al., 2024; Tu, Kim, et al., 2025). Informed consent was obtained from all participants, and the study received approval from the Human Studies Committee and Institutional Review Board at Washington University School of Medicine.

### Washington University 120 dataset

The detailed collection and preprocessing of the Washington University 120 (WU 120) dataset which has been described in prior research (Power et al., 2012, 2014, 2017). The study recruited 120 healthy young adults (aged 19-32 years, 60 females) to be scanned during a relaxed eyes-open fixation task. All participants were native English speakers and right-handed. They were recruited from the Washington University community and screened via a self-report questionnaire to ensure they had no current or past neurological or psychiatric conditions, as well as no head injuries resulting in a loss of consciousness for more than five minutes. Informed consent was obtained from all participants, and the study received approval from the Human Studies Committee and Institutional Review Board at Washington University School of Medicine.

Structural and functional MRI data were obtained using a Siemens MAGNETOM Trio Tim 3T Scanner (Erlangen, Germany) along with a Siemens 12-channel Head Matrix Coil. A T1-weighted sagittal magnetization-prepared rapid acquisition gradient-echo (MP-RAGE) structural image was captured with the following parameters: time echo (TE) of 3.08 ms, time repetition (TR) of 2.4 s, time to inversion (TI) of 1000 ms, a flip angle of 8°, and 176 slices with 1 × 1 × 1 mm voxel sizes. To ensure accurate alignment, an auto-align pulse sequence protocol provided by Siemens was used to position the functional scan slices parallel to the anterior commissure–posterior commissure plane of the MP-RAGE, aligning with the Talairach atlas (Talairach & Tournoux, 1988). During the functional MRI acquisition, participants were instructed to relax and focus on a black crosshair against a white background. The BOLD contrast-sensitive gradient-echo echo-planar imaging (EPI) sequence was employed, with a TE of 27 ms, a flip angle of 90°, and an in-plane resolution of 4 × 4 mm. Whole-brain EPI volumes, consisting of 32 contiguous 4-mm-thick axial slices, were collected every 2.5 seconds. Additionally, a T2-weighted turbo spin-echo structural image (TE = 84 ms, TR = 6.8 s, 32 slices with 1 × 1 × 4 mm voxels) was obtained in the same anatomical planes as the BOLD images to improve atlas alignment. Anterior→Posterior (AP) phase encoding was utilized for the fMRI acquisition, with the number of volumes collected from subjects ranging from 184 to 724 (mean = 336 frames, approximately 14.0 minutes).

Functional images were initially preprocessed through the following steps: (1) correction of intensity differences between odd and even slices caused by interleaved acquisition without gaps; (2) realignment to correct for head motion within and across functional runs; and (3) intensity normalization across runs, scaling each to a whole-brain mode value of 1000. Spatial normalization to a standard atlas space was performed for each participant using their corresponding MP-RAGE anatomical scan. Functional data were then resampled into a 3-mm isotropic voxel grid in Talairach space (Talairach & Tournoux, 1988), with motion correction and atlas transformation applied concurrently using a single cubic spline interpolation (Lancaster et al., 1995). The functional data went through additional custom artifact removal (Power et al., 2014) and were mapped to cortical surfaces (32k-fsLR).

### Baby Connectome Project dataset

Full-term infants (gestational age of 37-42 weeks) without any significant complications during pregnancy or delivery were recruited for the Baby Connectome Project (BCP) (Howell et al., 2019). All procedures were approved by the Institutional Review Boards of the University of North Carolina at Chapel Hill and the University of Minnesota. Informed consent was obtained from the parents of all participants. MRI images were acquired using a Siemens 3T Prisma scanner with a 32-channel head coil at both the University of Minnesota and the University of North Carolina at Chapel Hill while the infants were naturally asleep without the use of sedative medications. The imaging protocols included T1-weighted scans (TR=2400 ms, TE=2.24 ms, 0.8 mm isotropic; flip angle = 8°), T2-weighted scans (TR=3200 ms, TE=564 ms, 0.8 mm isotropic), spin echo field maps (SEFM) (TR=8000 ms, TE=66 ms, 2 mm isotropic, MB=1), and fMRI data (TR=800 ms, TE=37 ms, 2 mm isotropic, MB=8). For fMRI acquisition, a combination of Anterior→Posterior (AP) and Posterior→Anterior (PA) phase encoding directions was utilized, which were then combined into a single time series. An early subset of data was collected with a TR of 720 ms (N = 95). After conducting quality control and preprocessing using previously described methods including 30 parameter motion regression and motion censoring of volumes exceeding 0.2mm respiratory frequency notch-filtered FD (Kardan et al., 2022), the final cohort included 313 fMRI sessions from 181 individuals (95 females, aged 8 to 60 months, with a mean age of 19.1 months and a standard deviation of 8.3 months). The number of low-motion volumes obtained from participants ranged from 840 to 2100, with a mean of 1306 frames (approximately 16.9 minutes).

### Statistical analysis

In the following, we discuss three community detection methods: (1) WSBM, (2) Spectral Clustering (SC), and (3) K-means Clustering (Fig. 2.b). These algorithms were chosen because they can be applied to weighted FC matrices without thresholding and, in the implementations considered here, do not explicitly optimize an assortative community objective. Subsequently, we discussed the strategies for choosing the optimal number of communities for these three clustering algorithms (Fig. 2.c). The analyses code was written in MATLAB (version 2020b). The details of the iterations and tuning parameter specifications are mentioned in the Supplement. For the sections below, *n* and *K* denote the dimension of the FC matrix and number of communities respectively.

### Community detection algorithms

#### Weighted Stochastic Block Model

The WSBM is one of the most widely used community detection algorithms in machine learning and statistics (Aicher et al., 2013, 2015), particularly for identifying community structures within networks. Stochastic Block Model (SBM) and its weighted variant (WSBM) have wide applications in social (Ng et al., 2021; Tang & Yang, 2014), protein (Airoldi et al., 2008), and gene networks (Melo et al., 2024). In the WSBM, the nodes of the network are partitioned into *K* mutually exclusive and exhaustive communities, with each node assigned to exactly one of these *K* communities. Vertices within the same community exhibit similar connectivity patterns. In the case of fMRI data, we treat the brain regions of interest (ROIs) as vertices, and the FC matrix of the ROIs serves as the edge weights (Gordon et al., 2016; Seitzman et al., 2020). For each subject, the FC matrix is constructed using pairwise temporal correlation coefficients between ROI time series.

Mathematically, we denote the *n* × *n* FC-matrix as *A* for *n* brain ROIs with entries *A_i_*_j_ (edge weights) indicating the temporal correlation between time series ROI’s *i* and *j* and we do not apply Fisher *z*-transformation. We denote, *z_i_* as the membership indicator of *i^t^*^ℎ^ ROI and the corresponding probability of it belonging to a particular community (*k*). The community assignments are assumed to be random, independently and identically distributed (i.i.d.) with probability distribution *P*(*z_i_* = *k*) = *λ_k_*, for all 1 ≤ *k* ≤ *K* and 1 ≤ *i* ≤ *n*. Since the community assignments are mutually exclusive and exhaustive, ∑*_k_ λ_k_* = 1. Following (Aicher et al., 2015), we assume that the edge weights follow a normal distribution with community-specific parameters. A similar approach has been adopted in (Betzel, Bertolero, et al., 2018; Tooley et al., 2022). While the Fisher *z*-transformation stabilizes variance, it also equalizes variability across community interactions, thereby suppressing community-specific variability (Figs. S1.b, S1.c, S1.e, S1.f, S1.h, S1.i). In contrast, by avoiding the Fisher *z*-transformation, our approach preserves block-specific variability. The detailed description of the WSBM model can be found in the Supplement.

### Consensus partition algorithm for WSBM

Owing to the stochastic nature of the optimality criteria (Step 4 of the loglikelihood approach discussed in the Supplementary Materials), the community structure configurations might become heavily dependent on the optimal value of *K* after multiple algorithmic trials. Therefore, it is of interest to explore and compare the community structures obtained from WSBM in a more aggregated manner based on various community structure realizations. We performed consensus algorithm on the valid solutions from WSBM as a measure to evaluate the performance of the MLE solution among other valid solutions. We outline the algorithm for obtaining a consensus solution given a fixed community size. Since the community labels generated in two independent iterations may not align, we first rearrange the community labels using the Hamming distance (Hamming, 1950). Let *z* = (*z*_1_, *z*_2_, … , *z_n_*) and 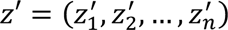 denote the estimated community labels from two independent iterations with community size *K*. Define, G*_K_* = {*π*: {1,2, … , *K*} ↦ {1,2, … , *K*}} as the set of all possible permutation of {1,2, … , *K*}. The Hamming distance between the community assignments *z* and *z*′ is given by

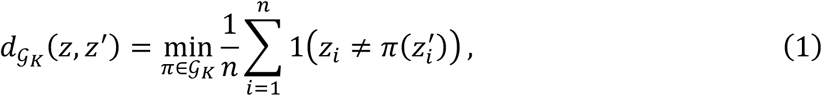

where 1 (*z_i_* ≠ π (*z_i_*′)) is the indicator function taking value 1 if (*z_i_* ≠ π (*z_i_*′) and 0 otherwise. In essence, the Hamming distance measures the minimum average proportion of mismatches between *z* and *z*′ across all possible community relabeling of *z*′. The minimization problem in equation (1) has a complexity of *O*(*n* + *K*^3^) using the Hungarian algorithm (Kuhn, 1955). The consensus assignment was then derived from the mode assignment for each node across the matched community labels. The consensus algorithm also allows us to analyze the community structures for different values of *K* which may serve as candidate community numbers, namely those for which the 95% confidence interval for the bootstrap log-likelihood difference includes zero, but are excluded by the decision rule in Step 4. Further details about the consensus algorithm and its implementation can be found in the Supplementary Materials.

#### Spectral Clustering

Spectral Clustering is a non-parametric clustering algorithm often used in machine learning (Hastie et al., 2009; Newman, 2006). The method has also been used to detect community structure in network data (Abbe, 2018; Rohe et al., 2011). Spectral Clustering is found to be useful non-linear structure of data as it tries to perform clustering on the eigenspace of a graph Laplacian constructed from a similarity matrix. The similarity matrix captures the pairwise similarity of data points based on a chosen distance metric. In our analysis, we used dissimilarity between pairs of ROIs as the individual Pearson correlation indices between the corresponding rows of the FC are subtracted from 1. This dissimilarity measure defines a weighted graph on the ROIs, and the associated Laplacian captures the network structure while its leading eigenvectors provide a low-dimensional embedding for clustering functional communities. The algorithm describing the Spectral Clustering can be found in the Supplementary Materials.

#### K-means Clustering

K-means Clustering is one of the most widely used clustering algorithms in machine learning, statistics, and various other disciplines (Duda et al., 2012; Hastie et al., 2009). K-means Clustering, known for its computational efficiency, has been used in PET and fMRI studies of the adult brain and has successfully recovered central and peripheral brain communities (Thirion et al., 2014; Tononi et al., 1998). Given the number of communities, the K-means Clustering algorithm identifies the *K* cluster centroids and assigns each data point to the community corresponding to the closest centroid. In the context of fMRI studies, we use correlation distance as the dissimilarity measure to determine the proximity of data points to the centroids. The detailed description of the K-means Clustering algorithm can be found in the Supplementary Materials.

#### Run time complexity of the algorithms

There is a trade-off in time complexity for these three algorithms. The following table (Table S2) in the Supplementary Materials shows the comparison of three algorithms in terms of runtime. Spectral Clustering requires an additional *O*(*n*^2^) time to store the dissimilarity matrix. But such computation is only needed once even for multiple independent iterations. Notably, Spectral Clustering and K-means Clustering are much faster than WSBM in dense networks.

### Benchmarking clustering algorithms

To benchmark the community assignments obtained with the three primary clustering algorithms, we compared the adult solutions with the Infomap-derived Gordon atlas and the infant solutions with the Infomap-derived Kardan atlas. We additionally applied Louvain modularity maximization to the BCP dataset only, providing an assortativity-based benchmark for the infant community structure.

#### Louvain Clustering

Louvain Clustering is a widely used community detection method for identifying modular structure in complex networks (Blondel et al., 2008). The method has also been extensively applied to functional brain networks, including networks containing both positive and negative connections (Rubinov & Sporns, 2010). Unlike K-means and Spectral Clustering, Louvain Clustering operates directly on the weighted network and does not require the number of communities to be specified in advance. Instead, communities are identified by approximately maximizing a modularity objective that measures the extent to which within-community connectivity differs from that expected under a corresponding null model. In our analysis, the ROIs were treated as nodes and the entries of the group-averaged FC matrix were used as signed edge weights. We employed a signed modularity formulation with symmetric treatment of positive and negative connections and examined community structure over a range of resolution parameters. The detailed description of the Louvain Clustering algorithm can be found in the Supplementary Materials.

#### Adult Infomap Clustering under the Gordon cortical atlas

The Gordon cortical atlas, introduced by (Gordon et al., 2016), is a functionally defined parcellation of the human cerebral cortex constructed from high quality resting state fMRI data (WU120). The atlas was generated by identifying boundaries where patterns of functional connectivity change sharply across the cortical surface, producing parcels that are internally homogeneous in their connectivity profiles and externally distinct from neighboring regions. Each parcel is assigned to one of several large-scale functional networks by applying Infomap to a thresholded FC matrix retaining 1–3% of all possible connections, excluding connections between parcels less than 20 mm apart, followed by literature-guided manual curation. Since it is grounded in functional connectivity rather than anatomy alone, the Gordon atlas is particularly well suited for studies of functional network architecture, individual differences, and brain behavior relationships, and it has been widely adopted in connectomics and systems neuroscience. To facilitate comparison across methods, we quantified the similarity between the network communities identified by the three clustering algorithms for the adult datasets (HCP and WU 120) and those defined by the Gordon atlas.

#### Infant Infomap Clustering the Kardan cortical atlas

The Kardan infant functional atlas (Kardan et al., 2022) is a functionally defined atlas specifically designed to characterize large scale brain networks in infants using fMRI data (BCP) acquired during early development, using the Infomap algorithm on thresholded sparse FC matrix and literature-guided experimenter curation. Prior work has demonstrated that the Gordon adult 333 areas have relatively high homogeneity with infant and toddler functional connectivity data (Tu, Myers, et al., 2025). Therefore, the 333 Gordon functional area parcels were used. However, research has demonstrated reduced long-range connectivity in higher order association cortex (Eggebrecht et al., 2017; Tu, Myers, et al., 2025). Recognizing that infant brains differ substantially from adult brains in anatomy, maturation, and functional connectivity, the Kardan atlas derived a functional network using the Infomap algorithm based on FC from infants and toddlers (8-26 months), rather than relying on adult based parcellations. FC matrices were thresholded at edge densities ranging from 1% to 10%, binarized, and used as inputs to the Infomap community detection algorithm. By capturing the immature but organized network structure of the infant brain, the Kardan infant atlas provides an essential framework for studying early functional network development, brain maturation trajectories, and the emergence of later cognitive and behavioral phenotypes, and is particularly well suited for infant connectomics. For the infant dataset (BCP), we compared the network communities identified by the three clustering algorithms with the Kardan atlas using similarity metrics. Importantly, while the participants used do not fully overlap between the current study and that from Kardan and colleagues, the same BCP data was used to derive communities, so the difference is mostly in methodology.

### Metrics to determine the optimal number of communities

To determine the optimal number of communities K, we considered the bootstrap log-likelihood-difference criterion for WSBM (Aicher et al., 2013, 2015; Faskowitz & Sporns, 2020), the silhouette index (Kaufman & Rousseeuw, 2009), and several post hoc measures (Sanchez-Rodriguez et al., 2021), including NMI, Hamming distance, VI distance (Meilă, 2007), modularity (Newman & Girvan, 2004), the Calinski–Harabasz index (Calinski & Harabasz, 1974), the C-index (Milligan & Cooper, 1985), the Dunn index (Dunn, 1973), and zRand (Akiki & Abdallah, 2019; Traud et al., 2011). Detailed definitions, selection rules, and implementation procedures are provided in the Supplementary Materials.

## Results

In this section, we present the performance of the three clustering algorithms on both synthetic and real-world datasets. Fig. 2 provides a diagrammatic overview of the three clustering methods and analyses in this study. To benchmark the empirical results, we compared the adult community assignments with the Infomap-derived Gordon atlas (Gordon et al., 2016) and the BCP community assignments with the Infomap-derived Kardan atlas (Kardan et al., 2022). For the BCP dataset, we also evaluated Louvain Clustering as an additional modularity-based benchmark. Because of the stochasticity of solutions from random initializations, throughout this manuscript, whenever we present an example solution of community assignment labels, we selected the solution with the highest log-likelihood among all valid solutions (WSBM), or the solution with the highest silhouette index among all 1000 replications (Spectral Clustering and K-means Clustering). In addition, we implement a consensus procedure for the WSBM by aggregating multiple candidate partitions to identify the community structure that is most stable and consistently supported across repeated realizations. Fig. S3 provides a diagrammatic representation of the consensus method implemented in our analyses. We restrict the use of the consensus approach to the WSBM because, for the other two algorithms, none of the post hoc criteria yielded an optimal choice for the number of communities *K*.

### Clustering results for synthetic data

To assess recovery of the ground-truth community number, we generated 100 independent synthetic FC matrices at each of *K* = 5, *K* = 7, and *K* = 10, using block-specific connectivity means and variances derived separately from the HCP and BCP datasets (Figs. 1 and S1). Each matrix was analyzed with all three clustering algorithms across a range of candidate community numbers. The heatmaps show the percentage of synthetic matrices for which each criterion selected each candidate value, with darker shades of blue indicating higher selection frequencies and the red-outlined columns denoting the ground-truth community number (Figs. 3.e and 3.f). Across all three community numbers and both connectivity parameterizations, the WSBM log-likelihood-difference criterion yielded the strongest and most consistent concentration of selections at the ground truth. This pattern was also evident in the representative simulation with *K* = 5, for which the bootstrap log-likelihood-difference criterion correctly selected the true community number (Fig. 3.a). Although several post hoc criteria recovered the ground truth in individual settings, their selections were generally more dispersed or shifted toward neighboring values, and their performance varied across clustering algorithms and connectivity parameterizations. Together, these results indicate that the WSBM log-likelihood-difference criterion provides a more reliable approach for recovering the underlying community number than the post hoc criteria considered here.

Beyond recovery of the community number, we examined the similarity and stability of the estimated community assignments. At the ground-truth values, WSBM, Spectral Clustering, and K-means Clustering recovered broadly comparable block structures under both the HCP and BCP informed simulation settings (Fig. S2). Spectral Clustering produced the most reproducible partitions across random initializations, with NMI values close to one and Hamming distances close to zero, whereas WSBM and K-means Clustering exhibited somewhat greater between-replication variability (Figs. 3.c and 3.d). Among the other post hoc metrics for selecting the optimal number of communities, modularity provided the most reliable recovery of the ground-truth community number. For WSBM, modularity consistently concentrated its selections at the true value of *K* under both the HCP- and BCP-informed settings; for Spectral Clustering and K-means Clustering, its performance was strongest when the true community number was *K* = 7 or *K* = 10, with greater dispersion when the true community number was *K* = 5 (Figs. 3.e and 3.f). The Calinski–Harabasz index showed a similar pattern, frequently recovering *K* = 7 and *K* = 10, but occasionally favoring a smaller community number when the ground truth was *K* = 5, particularly for K-means Clustering. When used to select *K*, the stability-based criteria showed pronounced algorithm-specific behavior. For WSBM, NMI, Hamming distance, and VI distance generally concentrated their selections at the ground truth, with some dispersion toward adjacent values. For Spectral Clustering, these criteria consistently favored *K* = 2, irrespective of the ground-truth community number or connectivity parameterization, indicating that highly reproducible partitions can nevertheless be substantially coarser than the underlying community structure. For K-means Clustering, the stability-based criteria performed inconsistently when the ground truth was *K* = 5, but more frequently recovered or closely approximated the true value when the ground truth was *K* = 7 or *K* = 10. The C-index produced comparatively dispersed selections, particularly for K-means Clustering, whereas zRand recovered *K* = 7 and *K* = 10 more reliably than *K* = 5. The Dunn index exhibited the weakest and most variable performance, often favoring smaller community numbers for ground truths of *K* = 7 and *K* = 10 under WSBM and Spectral Clustering, while favoring larger community numbers in several settings with a ground truth of *K* = 5 (Figs. 3.e and 3.f). These findings underscore that the reproducibility or apparent separation of an estimated partition does not necessarily imply accurate recovery of the underlying community number. The more consistent performance of the WSBM log-likelihood-difference criterion therefore supports its use as the primary approach for selecting *K* in the empirical analyses.

### Comparisons between methods applied to real data

#### Clustering results for adult datasets

We fit all the three algorithms to mean FC data from two young adult datasets (HCP and WU 120) across a range of *K* from 2 to 20, with 1000 replications for each value of K. For WSBM, due to algorithmic runtime complexity, we recovered less than 1000 valid clustering solutions for *K* > 4 (Table S6). Specifically, for *K* > 10, we recovered less than 200 valid clustering solutions, which has also been noted for rat cortex (Faskowitz & Sporns, 2020). Based on the difference log-likelihood plot and Step-4, using 2000 bootstrap iterations, we selected *K* = 11. The choice corresponds to the smallest value of *K* for which the 95% bootstrap confidence interval for the difference in log-likelihood includes the null value of zero (Figs. 4.a and 4.c). For all three algorithms, there was no clear maximum in the silhouette index for either the HCP or WU 120 adult datasets (Figs. 4.b and 4.d). To appreciate the spatial distribution of the communities, we also visualized the example solutions at *K* = 11 on the cortical surface plot (Figs. 5.a-5.f). For both the datasets, final community assignment solutions obtained using all three algorithms are in high agreement (peak NMI ∼ 0.7) with communities defined in the Gordon atlas (Gordon et al., 2016) (Figs. 5.g-5.i). For *K* = 11 and for HCP data, the median NMI values across 1000 replications for WSBM, Spectral Clustering and K-means Clustering were 0.63, 0.68 and 0.66 respectively. Similarly, for WU 120, the corresponding median NMI values were 0.67, 0.7 and 0.68 respectively.

**Figure 4.**
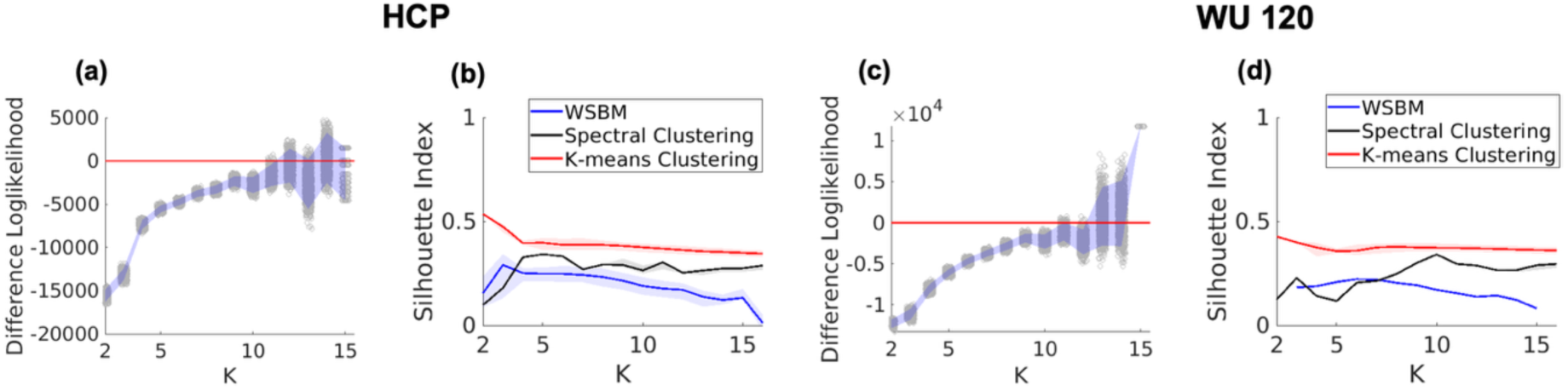
Detection of optimal number of communities for the adult datasets. **(a)** Bootstrap log-likelihood difference plot of the solutions from WSBM for HCP data. The red line corresponds to the null value zero and the shaded regions correspond to the 95% confidence interval for the bootstrap log-likelihood. **(b)** Silhouette index for HCP data. The solid lines denote median and shaded regions in denote the interquartile range values across 1000 replications. **(c)** Bootstrap log-likelihood difference plot of the solutions from WSBM for WU 120 data. The red line corresponds to the null value zero and the shaded regions correspond to the 95% confidence interval for the bootstrap log-likelihood. **(d)** Silhouette index for WU 120 data. The solid lines denote median and shaded regions in denote the interquartile range values across 1000 replications.

**Figure 5.**
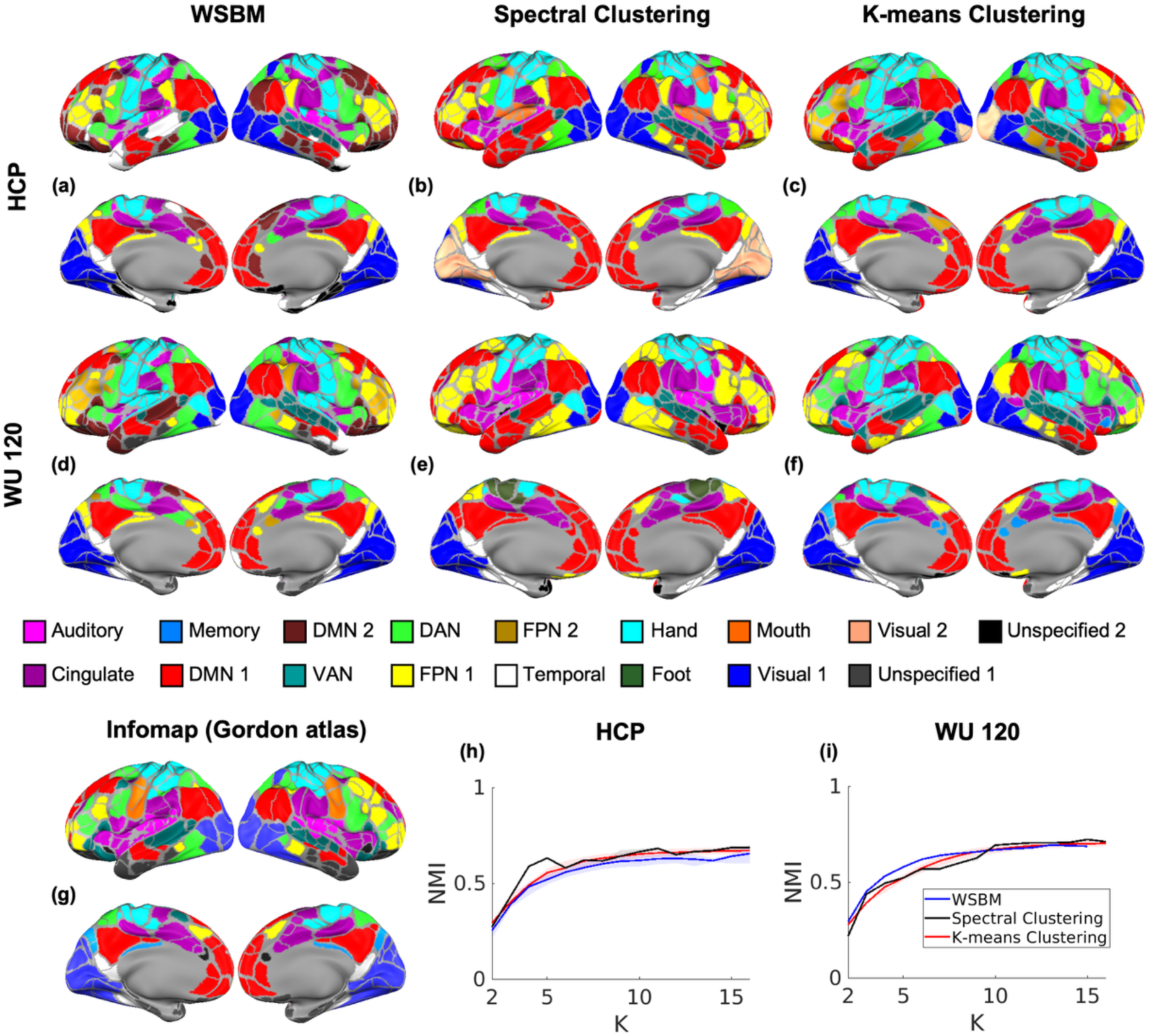
Community structure in adult FC and comparison with Gordon networks. **(a,d)** Brain surface plot of the solution with highest log-likelihood from WSBM for *K* = 11 for HCP and WU 120 data respectively. **(b,e)** Brain surface plot of the solution with highest silhouette index from Spectral Clustering for *K* = 11 for HCP and WU 120 data respectively. **(c,f)** Brain surface plot of the solution with silhouette index from K-means Clustering for *K* = 11 for HCP and WU 120 data respectively. **(g)** Gordon atlas. **(h,i)** NMI with community assignment in the Gordon atlas for HCP and WU 120 respectively. The solid lines denote median and shaded regions in denote the interquartile range values across 1000 replications. DMN - Default Mode Network, VAN - Ventral Attention, DAN - Dorsal Attention, FPN – Frontoparietal, Hand – Somatomotor Hand, Foot – Somatomotor Foot, Mouth – Somatomotor Mouth.

To assess between-algorithm agreement, we calculated NMI across solutions at each level of *K* pairwise across three algorithms. K-means and Spectral Clustering showed strong agreement with each other having highest NMI values as 0.74 and 0.72 for HCP and WU 120 respectively, while WSBM exhibited moderate agreement with K-means and Spectral Clustering (NMI <0.4; Figs. 6.a and 6.b). Similarly, Hamming distance values were generally low (0.3-0.5) across most *K* values, indicating strong agreement among the three clustering algorithms (Figs. 6.c and 6.d). We also used several post hoc metrics discussed in the Supplementary Materials to estimate the number of communities for the two adult datasets. However, none of the post hoc metrics identified a clear global optimum (Figs. 6.g-6.l and 6.o-6.t).

**Figure 6.**
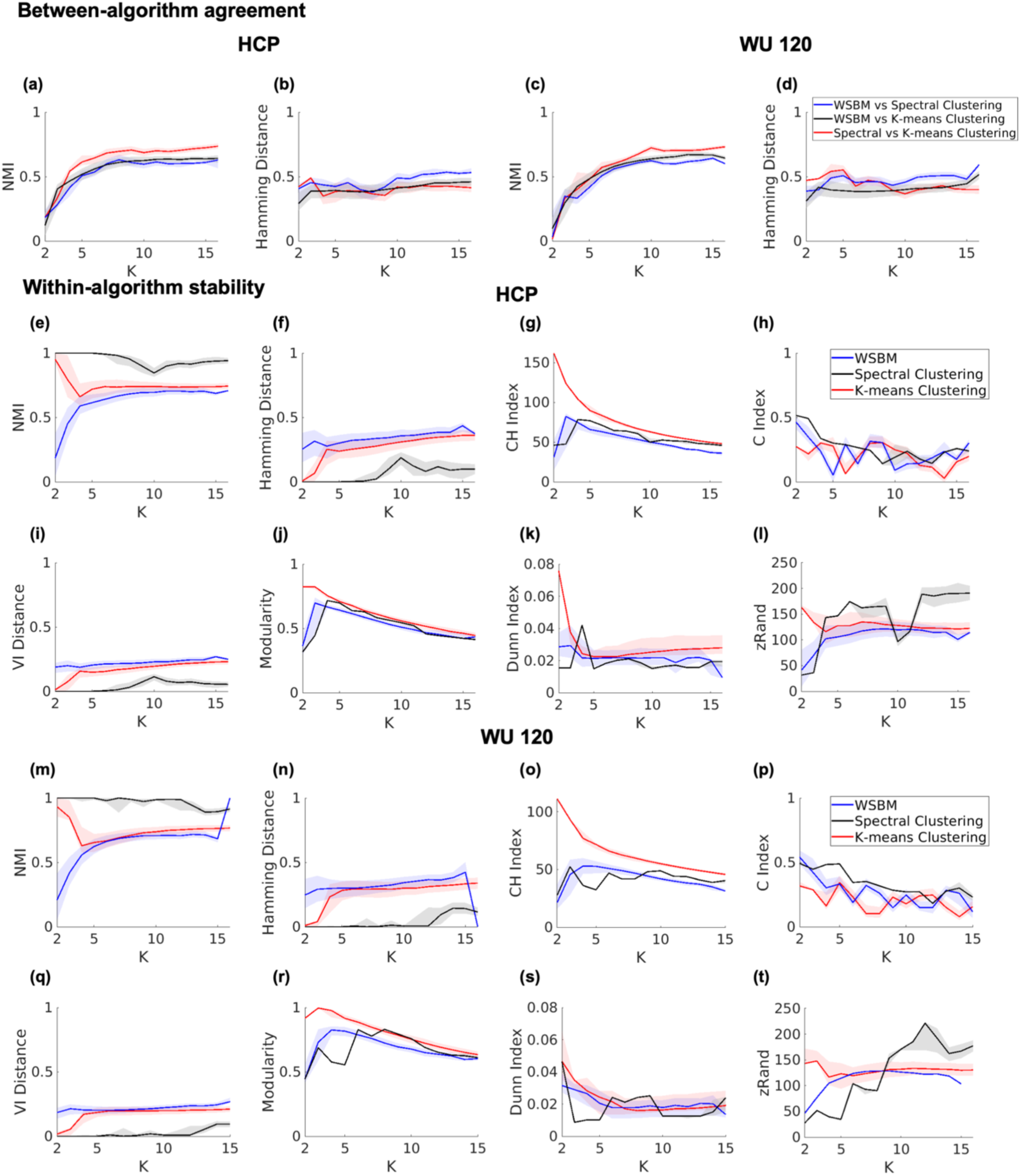
Between-algorithm community assignment agreement and within-algorithm community assignment stability. **(a,c**) NMI and Hamming distance for all pairwise community assignments between different clustering algorithms on HCP data. **(b,d**) NMI and Hamming distance for all pairwise community assignments between different clustering algorithms on WU 120 data. **(e-l)** Evaluation of different measures of clustering solutions for HCP data. **(e)** NMI, **(f)** Hamming Distance, **(g)** Calinski–Harabasz (CH) index, **(h)** C-index, **(i)** VI Distance, **(j)** Modularity, **(k)** Dunn Index, **(l)** zRand. The shaded regions represent the inter quartile ranges. **(m-t)** Evaluation of different measures of clustering solutions for WU 120 data. **(m)** NMI, **(n)** Hamming Distance, **(o)** Calinski–Harabasz (CH) index, **(p)** C-index, **(q)** VI Distance, **(r)** Modularity, **(s)** Dunn Index, **(t)** zRand. The shaded regions represent the inter quartile ranges.

To assess the stability across valid solutions within each algorithm, we calculated the pairwise NMI across 1000 solutions (or all valid solutions, if it were WSBM). For both young adult datasets, Spectral Clustering yielded the most stable solutions across all levels of *K* (NMI > 0.8). In contrast, K-means Clustering solutions had high stability at *K* = 2, but decreased to a plateau (NMI ∼0.65), while WSBM solutions started with low stability at *K* = 2, but increased to a plateau (NMI ∼0.65) when the number of communities exceeded 5 (Figs. 6.e and 6.m). Hamming distance followed a similar pattern, with lowest values in Spectral Clustering (between 0 and 0.2), while K-means Clustering and WSBM had moderately low values (between 0.2 and 0.4) (Figs. 6.f and 6.n).

Overall, all three clustering algorithms successfully recovered the common sensory and association networks that were agreed upon by the field based on resting-state and task fMRI data (Uddin et al., 2019). Solutions across algorithms were moderately similar at *K* = 11 (a.k.a. the optimal number of communities based on log-likelihood difference in WSBM), as evident in the quantitative measure of NMI and hamming distance, as well as the example solution at *K* = 11 visualized on brain surfaces (Figs. 5.b, 5.c, 5.e, 5.f, S5.b, S5.c, S5.e and S5.f) and on FC matrices sorted by the community labels (Figs. S4.a–S4.f present the FC matrices). Additionally, WSBM was able to identify Default Mode Network (DMN) 2 and Temporal Cingulo Opercular as peripheral regions relative to DMN1 for *K* = 11 (Fig. S10.a).

#### Clustering results for infant dataset

We fit all the three primary algorithms and Louvain Clustering to mean FC in the BCP dataset. WSBM, Spectral Clustering and K-means Clustering are preformed across a range of K between 2 to 20 and for 1000 replications. For Louvain clustering, we vary the resolution parameter from 0 to 4 in increments of 0.05, performing 1000 replications for each parameter value. For WSBM, due to algorithmic runtime complexity, we recovered less than 1000 valid clustering solutions for *K* > 6 (Table S6). For the BCP dataset, the difference in log-likelihood plot, derived from 2000 bootstrap iterations, suggested that the optimal number of communities is *K* = 15 (Fig. 7.a). In contrast, akin to the adult datasets, a distinguishable global maximum could not be identified using Silhouette index for any of the four algorithms (Fig. 7.b). Similar solutions were observed across the community detection methods (Figs. 7.c-7.f). While WSBM was able to identify communities in the somatomotor network organized by topography (a.k.a. hand, foot, and mouth), motor foot was clustered into motor hand in Spectral Clustering while clustered into cingulo-opercular network in K-means Clustering. Like WSBM, the Louvain algorithm also preserves the distinct hand, foot and mouth communities at *K* = 15. The Louvain analysis favored a *K* = 5 community solution, in contrast to the *K* = 15 solution selected by the WSBM log-likelihood-difference criterion (Figs. 7.a and 7.b). In the surface-based comparison at *K* = 15, Louvain produced a coarser somatomotor organization than WSBM, with the foot and mouth communities largely subsumed within the hand community (Figs. 7.c and 7.f). Communities including the DMN and Fronto-Parietal Network (FPN) were consistently split into anterior and posterior components of their adult network counterparts across all four methods. For *K* = 15 (Fig. 7.h), the NMI values for the BCP data were 0.67, 0.72, 0.70 and 0.67 for WSBM, Spectral Clustering, K-means Clustering and Louvain algorithm respectively, when compared to the Kardan network atlas (Kardan et al., 2022) (Fig. 7.g). Figs. S4.g-S4.i show the FC matrix plots and Figs. S5.g-S5.i show the community assignment labels visualized on brain surfaces.

**Figure 7.**
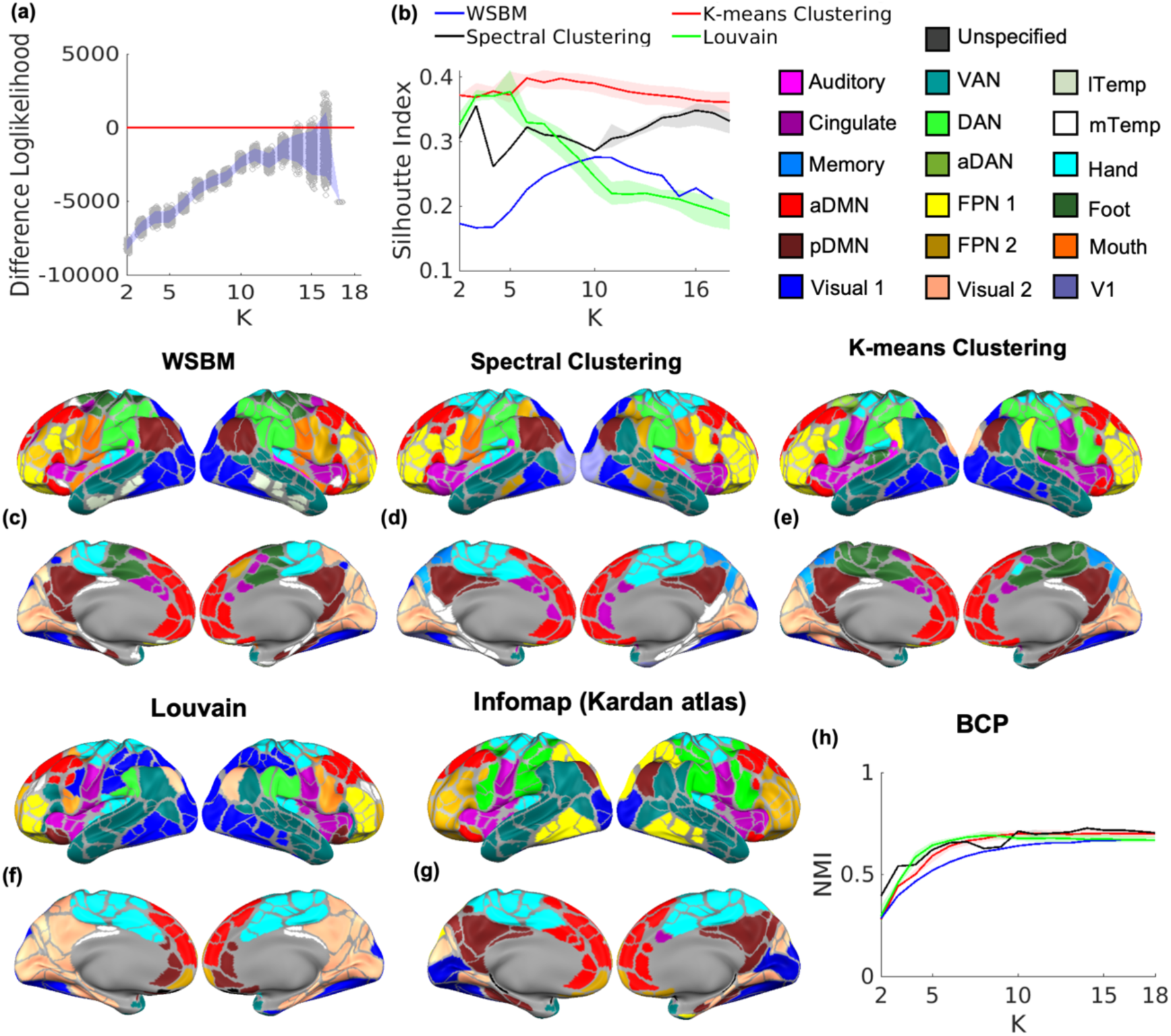
Detection of optimal number of communities for the infant dataset. **(a)** Bootstrap log-likelihood difference plot of the solutions from WSBM for BCP data. The red line corresponds to the null value zero and the shaded regions correspond to the 95% confidence interval for the bootstrap log-likelihood. **(b)** Silhouette index values with interquartile range for BCP data. **(c)** Brain surface plot of the solution with highest log-likelihood from WSBM for *K* = 15 for BCP data. **(d)** Brain surface plot of the solution with highest silhouette index from Spectral Clustering for *K* = 15 for BCP data. **(e)** Brain surface plot of the solution with silhouette index from K-means Clustering for *K* = 15 for BCP data. **(f)** Louvain solution for *K* = 15 for BCP data. **(g)** Kardan atlas. **(h)** NMI values with interquartile range between the community assignment solutions from four methods and Kardan atlas for BCP data. aDMN - Anterior Default Mode Network, pDMN - Posterior Default Mode Network, VAN – Ventral Attention, DAN – Dorsal Attention, FPN – Frontoparietal, lTemp - Lateral Temporal, mTemp - Medial Temporal, Hand - Somatomotor Hand, Foot - Somatomotor Foot, Mouth - Somatomotor Mouth, V1 – Primary Visual.

Within-algorithm stability differed across methods in the BCP dataset. Spectral Clustering produced the most reproducible partitions, with NMI values close to one and Hamming and VI distances close to zero across much of the examined range of *K*; Louvain and K-means Clustering showed intermediate stability, whereas WSBM stability increased progressively with *K* (Figs. 8.a, 8.b, and 8.e). The remaining post hoc indices exhibited method-dependent patterns and did not identify a consistent optimal number of communities across the four methods (Figs. 8.c, 8.d, and 8.f–8.h).

**Figure 8.**
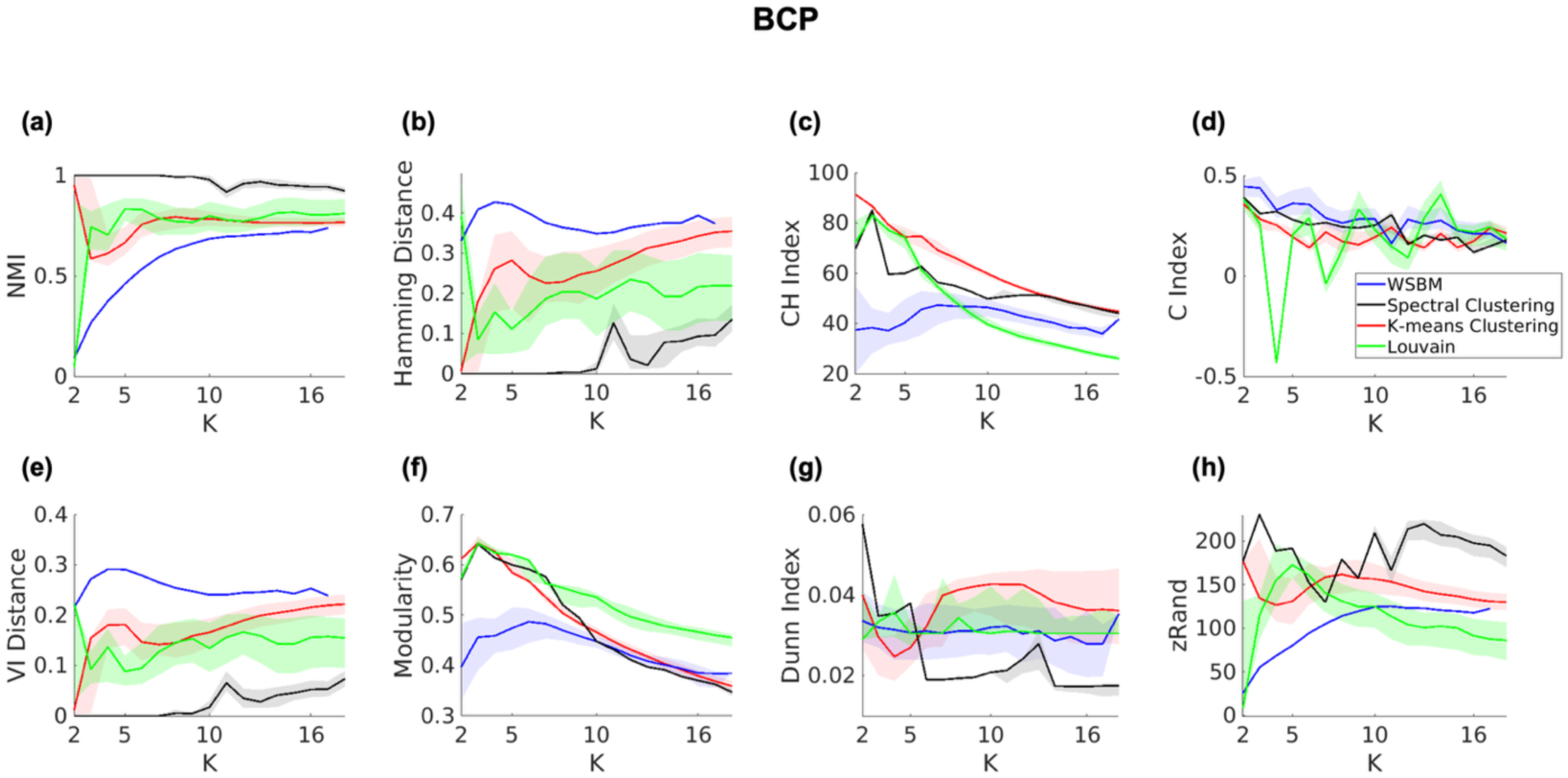
Evaluation of different measures of clustering solutions for BCP data. **(a)** NMI, **(b)** Hamming Distance, **(c)** Calinski–Harabasz (CH) index, **(d)** C-index, **(e)** VI Distance, **(f)** Modularity, **(g)** Dunn Index **(h)** zRand. The shaded regions represent the inter quartile ranges.

#### Consensus solutions for all datasets

Although we used the minimum value of *K* as the main criterion for selecting the optimal number of communities, the bootstrapped log-likelihood differences (Figs. 4.a, 4.c and 7.a) indicated that alternative choices of *K* were also plausible. Therefore, we evaluated consensus communities for all values of *K* where the null value 0 fell within the range of the bootstrap log-likelihood differences. To construct the consensus, we applied the consensus procedure described previously in the section on Statistical Analysis to all valid WSBM solutions for each fixed *K*. As reference communities, we chose the solution with the highest log-likelihood for that *K*. The brain surface plots of the consensus solutions for the HCP, WU 120, and BCP datasets are shown in Figs. S6.a–S6.d, S7.a–S7.c, and S8.a–S8.d, respectively. When comparing the consensus solutions (Figs. S6.a and S7.a) with the highest log-likelihood solutions for adults (Figs. 5.a and 5.d), the spatial distribution of networks were largely consistent with that at *K* = 11. In contrast, for the infant dataset (Fig. 7.c), the consensus solution (Fig. S8.a) diverged from the *K* = 15 solution only for one network, with the posterior DMN and Ventral Attention Network (VAN) combined into a single network. The NMI values between the MLE and consensus solutions are 0.85, 0.91 and 0.84 for HCP, WU 120 and BCP datasets respectively. These high values indicate very strong agreement between the community structure recovered by the MLE solutions and that identified by the consensus algorithm.

For each fixed *K*, we also plotted the rearranged community labels across independent replications, grouping them by community labels for the HCP, WU 120, and BCP datasets (Figs. S.6e–S6.h, S7.d–S7.f, S8.e–S8.h). These findings underscore both the reproducibility of the consensus procedure and its ability to yield interpretable community structures under multiple candidate values of *K*. For *K* = 14, the consensus algorithm identified Visual 2 as a peripheral region relative to Visual 1 (Fig. S10.d), suggesting that within-network connectivity is stronger in Visual 1 than in Visual 2 for the HCP data.

Finally, we examined the FC matrices averaged across subjects to assess community-level connectivity strength. The averaged FC matrices, grouped by community labels, are shown in Figs. S9.a–S9.k revealing mostly assortative structures high within community connections in the human brain (Table S3-S5). These patterns are consistent with those observed in Figs. S4.a–S4.f for the highest log-likelihood and silhouette index solutions.

## Discussion

In this article, we addressed the problem of determining the optimal number of communities in human FC data using the WSBM log-likelihood-difference criterion. Several important insights emerged from our results. First, the performance of commonly used cluster-quality indices varied across algorithms, ground-truth community numbers, and connectivity parameterizations. Modularity and the Calinski–Harabasz index frequently recovered the ground truth, whereas the remaining post hoc criteria did not provide consistent guidance across all simulation settings. Second, solution stability varied across algorithms in both the synthetic and empirical datasets. Spectral Clustering generally produced the most reproducible partitions, whereas WSBM and K-means Clustering showed greater variability across replications; between-method agreement was strongest between Spectral and K-means Clustering and more moderate for comparisons involving WSBM. At the selected values of *K*, the methods produced predominantly assortative communities that shared similarities with previously published communities from the same adult and infant datasets. Although the methods recovered a broadly shared set of major neural systems at matched values of *K*, they differed in the granularity and boundaries of the resulting communities. The predominantly assortative organization supports the biological relevance of modular structure and suggests that assortative community-detection methods can provide useful approximations for these data. Nevertheless, topology-flexible approaches remain valuable because they allow the observed assortative organization to emerge from the data rather than imposing it a priori, and the convergence across methods does not imply that they will select the same community granularity. Moreover, the communities recovered at the selected values of *K* were consistent with the known organization of neural systems described in the neuroscience literature.

Our findings are complementary to Brooks and colleagues, who showed that the performance and agreement of community-detection methods depend on the underlying network topology (Brooks et al., 2024). Whereas their study examined method dependence across heterogeneous topological regimes, our analysis focuses on selecting Kand evaluating the stability of the resulting partitions in dense, weighted adult and early-life functional connectomes. Together, these studies emphasize the importance of assessing robustness with respect to both the detection method and the selected community granularity.

The task of determining the number of communities in human connectomes has been explored in a number of studies. Previously, Yeo and colleagues fit a von Mises–Fisher model and conducted an instability analysis to select the number of communities, identifying *K* = 7, 10, 12, and 17 as plausible solutions, while ultimately recommending the boundary values *K* = 7 and *K* = 17 in an agnostic manner (Yeo, et al., 2011). Building on this framework, Tooley and colleagues applied a WSBM and adopted a similar instability-based rationale for selecting the number of communities (Tooley et al., 2022). Based on their instability analysis plot, they identified *K* = 7 as the solution for adolescent data, consistent with the *K* = 7 solution suggested by Yeo and colleagues.

However, relying on instability plots and local minima yields multiple plausible values of *K*, without a clear, principled basis for selecting one. Using a bootstrap-based log-likelihood difference approach, we present a principled way of choosing the optimal number of communities for human FC data. For each choice of *K*, we used the set of valid solutions to construct 95% bootstrap confidence intervals for the log-likelihood differences for *K* and *K* + 1. The optimal number of communities was chosen to be the smallest value of *K* for which this confidence interval contained the null value zero. Our results using the optimal *K* communities for adult and infant FC show strong agreement with previously reported parcellations (Gordon et al., 2016; Kardan et al., 2022), lending empirical support to our proposed guideline for selecting the optimal number of communities.

Our analysis using WSBM identified 11 communities across two independent young adult cohorts from the HCP and WU 120 adult datasets, adding credence to the conclusion that 11 can be treated as the optimal number of resting-state networks in adults. Notably, all three clustering algorithms exhibited similar community solutions both across replications and across algorithms, especially across Spectral and K-means Clustering. Further, the optimal *K* aligned closely with relatively high NMI values with the Gordon atlas (Gordon et al., 2016). However, some differences between algorithms in the derived communities were observed. For *K* = 11, WSBM identified one Visual network consistent with (Gordon et al., 2016) while the other K-means and Spectral Clustering split the network into Lateral and Medial Visual consistent with (Gordon et al., 2017).Furthermore, WSBM identified two distinct DMN subnetworks while K-means and Spectral Clustering only returned a single DMN network. While prior work has predominantly described the DMN as a single network (Gordon et al., 2016; Yeo, et al., 2011), our analysis with WSBM is consistent with prior evidence that the default mode network can be meaningfully subdivided into two distinct regions with different network topologies (Gordon et al., 2020).

For the infant brain, WSBM was able to identify 15 communities. At matched values of *K*, the three primary methods produced broadly comparable community solutions; however, Louvain Clustering favored *K* = 5, in contrast to the *K* = 15 solution selected by the WSBM log-likelihood-difference criterion. Analysis on the BCP data suggested that infant brain communities differ from adult brain communities in terms of number of detected communities and spatial topology. In the surface-based comparison at *K* = 15, Louvain recovered the broad functional scaffold but produced a coarser somatomotor organization, with the foot and mouth communities largely subsumed within the hand community. Additionally, for the infant brain, WSBM, K-means, and Spectral Clustering all demonstrated the DMN and FPN are divided into two distinct communities with anterior and posterior components, consistent with prior work in infants using Infomap clustering (Eggebrecht et al., 2017; Kardan et al., 2022; Tu, Myers, et al., 2025). Consensus results also confirmed that community assignments remain consistent across different choices of *K*. Optimal community assignments also have moderately high NMI values with previously published the Infomap solutions in the infant age group (Kardan et al., 2022). While Infomap solutions for neonate and toddler datasets can be used to identify more ‘adult like’ networks by increasing the density of the graph (Sylvester et al., 2023; Tu, Myers, et al., 2025; Tu, Wang, et al., 2025), these long-range connections are weakly connected at these ages. The latter findings may be explained by early stages of brain development or by the fact that the scans were acquired during sleep. Together, these findings indicate that, despite the presence of an early adult-like functional scaffold, the infant brain exhibits developmentally distinct mesoscale organization, with the WSBM-selected solution separating several higher-order association networks into anterior and posterior components.

### Limitations

This article provides some insight into selecting the optimal number of communities for adult and infant brain connectomes. Our analysis suggests that post hoc community-evaluation metrics do not perform uniformly across clustering algorithms, community numbers, and connectivity parameterizations, even though some criteria, particularly modularity and the Calinski–Harabasz index, performed well in individual settings. Here we demonstrate that WSBM, with the help of difference log-likelihood, is an effective method to obtain the unknown number of communities. Furthermore, we illustrate the fact that the community structure of the infant brain is different from that of the adult brain. However, one needs to keep in mind that some of the community detection algorithms mentioned above (including WSBM) are affected by detectability issues and resolution limits, which can impact their performance. The detectability limit is known to depend on the graph’s characteristics rather than the specific algorithm used (Reichardt & Leone, 2008). Additionally, the WSBM is also subjected to a resolution limit. For a network of size *n*, the maximum number of detectable communities is √*n* when using maximum log-likelihood estimation (Choi et al., 2012; Peixoto, 2013).

As discussed earlier, the community assignments in WSBM are based on the maximum posterior assignment probability. However, this does not automatically guarantee that the number of estimated communities will match the desired number specified as input (Faskowitz & Sporns, 2020). This leads to an important limitation of our bootstrapped samples, where our calculated bootstrapped difference in log-likelihood samples from increasingly smaller samples with higher numbers of *K*. Despite that, a few lines of evidence add credence to our identified optimal number of communities. First, the failure to recover valid solutions at higher community number *K* could also be suggestive in itself of a sub-optimal *K* and that the optimal community count is at a lower *K*. Secondly, the consensus from solutions with higher *K* choices converge to the solution with our optimal solution by the difference in log likelihood method. Third, at the selected value of *K*, we observed reproducible within-method solutions and moderate agreement across methods, although the degree of stability varied across algorithms. Moreover, the communities recovered from the optimal *K* choice are consistent with the known neural systems organization from decades of anatomical and functional evidence in human neuroscience studies. Finally, from a computational standpoint, while these methods are scalable to higher-resolution networks, their scalability can vary significantly depending on the algorithm.

Another important limitation is that we conducted our community detection algorithm on one single mean FC matrix across the whole population, whether our optimal number of communities is stable with regards to different sub-samples of the population is an outstanding question (Yeo, et al., 2011). Other recent studies also highlighted the potential caveats of smoothing out individual differences when community detection was conducted on average data, especially if the spatial representation of communities varies across individuals in the population (Gordon et al., 2017; Hermosillo et al., 2024; Tu et al., 2026). Additionally, we have assumed a non-overlapping community organization (a.k.a. hard parcellation), while others have attempted to describe the human connectome using a “soft parcellation” (Bryant et al., 2026; Faskowitz et al., 2020; Harrison et al., 2015; Lin et al., 2018; Najafi et al., 2016; Tu et al., 2026).

## Supporting information

Supporting information for this article is available in the Supplementary Material. Human Connectome Project data is available at https://db.humanconnectome.org/. WashU 120 data is available at https://openneuro.org/datasets/ds000243/versions/00001. Baby Connectome data is available at https://nda.nih.gov/edit_collection.html?id=2848. The code for reproducibility of these results can be found in https://github.com/ayoushmanb/BrainNetworks_WSBM_SC_KM.

## Author contributions

Ayoushman Bhattacharya: Formal analysis; Investigation; Methodology; Software; Validation; Visualization; Writing – original draft; Writing – review & editing. Nilanjan Chakraborty: Conceptualization; Formal analysis; Investigation; Methodology; Project administration; Validation; Visualization; Writing – original draft; Writing – review & editing; Supervision. Jiaxin Cindy Tu: Data curation; Methodology; Software; Resources; Writing – review & editing. Xintian Wang: Data curation; Visualization; Writing – review & editing. Donna Dierker: Data curation; Resources; Writing – review & editing. Andy Eck: Software; Writing – review & editing. Jed T. Elison: Resources; Writing – review & editing. Soumen Lahiri: Supervision; Validation; Writing – review & editing. Adam Eggebrecht: Software; Writing – review & editing. Muriah D. Wheelock: Conceptualization; Data curation; Formal analysis; Funding acquisition; Investigation; Methodology; Project administration; Resources; Software; Supervision; Validation; Visualization; Writing – original draft; Writing – review & editing.

