## supplement for "Comparative Evaluation of Assumption Lean Community Detection Methods for Human Connectome Networks"

### Description of clustering algorithms

#### Weighted Stochastic Block Model

Following the notations in the main manuscript, the likelihood of the FC matrix $A$ conditioned on the parameters can be written as follows:

$$P(A|z,\mu,\Sigma) = \prod_{1\le i<j\le n} N\left(A_{ij}\middle|\mu_{z_iz_j},\sigma^2_{z_iz_j}\right), \quad \text{(S1)}$$

where $N\left(A_{ij}\middle|\mu_{z_iz_j},\sigma^2_{z_iz_j}\right)$ denotes the density of a normal distribution with mean $\mu_{z_iz_j} \in \mathbb{R}$ and connectivity variance $\sigma_{z_iz_j} > 0$. The matrix $\mu_{K\times K} = (\mu_{ab})$ is the mean connectivity of the group interaction and $\Sigma_{K\times K} = (\sigma^2_{ab})$ is the variability in connection of the group interaction. We assume that the weighted adjacency matrix $A$ is symmetric, i.e., $A_{ij} = A_{ji}$ for all $1 \le i < j \le n$ and $A_{ii} = 1$ which we disregard in our analysis since there is no randomness involved in estimating the diagonal elements. We fit $z_i$ with flat prior $\pi(z_i) = \frac{1}{k}$ for all $1 \le i \le n$. For the canonical parameter vector, $\eta = (\eta_{ab})^T{}_{1\le a,b\le k}$ with $\eta_{ab} = \left(\frac{\mu_{ab}}{\sigma^2_{ab}}, -\frac{1}{2\sigma^2_{ab}}, -\frac{\mu^2_{ab}}{\sigma^2_{ab}}\right)^T$, we assume the conjugate prior from an exponential family taking the form,

$$\pi(\eta) = \frac{1}{Z(\tau)} \prod_{1\le a,b\le K} \exp\{\eta^T\tau\},$$

where $\tau$ is the vector for hyper-parameters and $Z(\tau)$ is the normalized constant. Assuming the independence of the priors, we finally have the joint prior as follows,

$$\pi(z,\eta) = \prod_{1\le i\le n} \pi(z_i)\ \frac{1}{Z(\tau)} \prod_{1\le a,b\le k} \exp\{\eta^T\tau\}.$$

Therefore, the log-likelihood of the joint pdf can be written as

$$l(A,z,\eta) = \log P(A|z,\mu,\Sigma) + \log \pi(z,\eta).$$

##### Estimation of the model parameters for WSBM

A straightforward maximization of (S4) is not feasible because the community memberships are unknown. To address this, we use the Variational Bayes technique (Aicher et al., 2013) to estimate the community memberships $(z_i)$ and the community-specific parameters $(\mu,\Sigma)$. The goal is to approximate the joint posterior distribution of $(z,\mu,\Sigma)$ by minimizing the Kullback-Leibler (KL) divergence between the true posterior distribution and the product of the marginal posterior distributions of $z$ and $(\mu,\Sigma)$.

We implemented the WSBM using the publicly available MATLAB implementation (https://aaronclauset.github.io/wsbm/) with the following parameter settings: '`algType`' = '`vb`', ‘alpha’ = 0, ‘networkType’ = ‘sym’, ‘parallel’ = 0, ‘E_Distr’= ‘None’, ‘W_Distr’ = ‘Normal’, ‘numTrials’ = 1, ‘mainMaxIter' = 80, ‘mainTol’ = 0.001, ‘muMaxIter’ = 50, ‘muTol’ = 0.001, ‘mexFile’ = 1.

##### Consensus for WSBM

In this section, we describe the consensus algorithm for WSBM as follows.

- Step 1: For a fixed $K$, select a community assignment arbitrarily from the set of all valid solutions to serve as the reference label. In our analysis, the solution corresponding to the maximum log-likelihood was be chosen for the reference.
- Step 2: Compute the Hamming distance between the reference label and each of the remaining solutions.
- Step 3: For each solution, determine the permutation mapping $\pi$ that minimizes the optimization problem described in equation (1). The communities are relabeled accordingly using the minimizer $\pi$.
- Step 4: Finally, each node is assigned to the community obtained as the minimizer in Step 3 appearing most frequently across the relabeled solutions and thus forming the consensus community assignment.

The overview of the algorithm is presented in Fig. S3. This algorithm resolves label-switching ambiguities, resulting in a stable and interpretable community structure. Since relabeled community assignments obtained in Step 3 are equivalent to the original community assignment up to a permutation of class labels, the number of communities for each assignment remains unchanged.

#### Spectral Clustering

We define the dissimilarity measure $d_{ij} = 1 - r_{ij}$, where $r_{ij}$ is the Pearson correlation index between $i^{th}$ and $j^{th}$ rows of the FC matrix. This transformation converts $r_{ij}$’s to a non-negative dissimilarity measure $d_{ij}$’s. The similarity matrix $S_{n\times n} = (S)_{1\le i,j\le n}$ is constructed as $S_{ij} = \exp(-d_{ij}^2)$ denoting the similarity between ROIs $i$ and $j$. Thus, we formally define the graph Laplacian matrix as $L = D - S$, where $D$ is a diagonal matrix with entries $D_{ii} = \sum_{1\le j\le n; j\ne i} S_{ij}$. The Spectral Clustering algorithm is described as below.

- Step 1: Find the eigenvectors $X_1, \ldots, X_K \in \mathbb{R}^n$ corresponding to the $K$ eigenvalues of $L$ that are smallest in terms of their absolute values.
- Step 2: Form the matrix $X = [X_1, \ldots, X_K] \in \mathbb{R}^{n\times K}$ by putting the eigenvectors into the columns.
- Step 3: Treating each of the $n$ rows in $X$ as a point in $\mathbb{R}^K$ and run the K-means algorithm with $K$ clusters, which creates $K$ nonoverlapping sets $C_1, \ldots, C_K$ whose union is $1, \ldots, n$.
- Step 4: Return the communities $C_1, \ldots, C_K$. This means that ROI $i$ is assigned to the cluster $g$ if the $i^{th}$ row of $X$ is assigned to $C_g$ in step 3.

We used the built-in Matlab function '`spectralclutser`' using the following parameter settings: '`Distance`' = '`correlation`', '`LaplacianNormalization`' = '`None`', '`ClusterMethod`' = '`kmeans`', '`MaxIter`' = 1000, '`Replicates`' = 1, '`TolFun`' = 0.0001.

After detecting the optimal $K$, the final solution is chosen to be the one with maximum silhouette index for that value of $K$. The K-means Clustering algorithm was implemented using the built-in MATLAB function '`kmeans`' (https://www.mathworks.com/help/stats/kmeans.html). Spectral Clustering was implemented using the built-in MATLAB function '`spectralcluster`' (https://www.mathworks.com/help/stats/spectralcluster.html). Additional details on the parameter choices for the algorithms are provided in the Supplementary Material.

#### K-means Clustering

Let $y_i$ denote the $i^{th}$ row of the weighted adjacency matrix $A$ for $1 \leq i \leq n$. Thus, each $y_i$ is a $n$-dimensional vector. As defined in previous section, we used $d_{ij} = 1 - r_{ij}$ as the correlation index-based dissimilarity measure to the distance of the data points from the centroids. The K-means Clustering algorithm is described in steps as follows:

- Step 1: Randomly assign $K$ centroids: $\mu_1, \dots, \mu_K$.
- Step 2: Classify $n$ data points $y_1, \dots, y_n$ by the closest mean.
- Step 3: Recompute the cluster centroids based on the cluster formed in step 2.
- Step 4: Repeat steps 2-3 until there is no change in the centroids.

We used the built-in Matlab function '`kmeans`' using the following parameter settings: '`MaxIter`' = 1000, '`start`' = '`plus`', '`Distance`' = '`correlation`', '`Replicates`' = 1, '`TolFun`' = 0.0001.

#### Louvain Clustering

Let $A = (A_{ij})$ denote the weighted adjacency matrix. To account for the signed nature of the functional connectivity network, we separately define the positive and negative parts of the adjacency matrix as $A_{ij}^{+} = \max(A_{ij}, 0)$ and $A_{ij}^{-} = \min(A_{ij}, 0)$. Define the corresponding positive and negative node strengths by $s_i^{+} = \sum_j A_{ij}^{+}$, $s_i^{+} = \sum_j A_{ij}^{-}$ respectively and the total positive and negative network weights by $v^{+} = \sum_{ij} A_{ij}^{+}$, $v^{-} = \sum_{ij} A_{ij}^{-}$ respectively. For a partition of the ROIs into communities, let $z_i$ denote the community assignment of $i^{th}$ ROI. Following the symmetric treatment of negative weights implemented in the Brain Connectivity Toolbox (Rubinov & Sporns, 2010), the modularity of the partition is defined as

$$Q = \frac{1}{v^{+} + v^{-}} \sum_{ij} \left[ \left( A_{ij}^{+} - \gamma \frac{s_i^{+} s_j^{+}}{v^{+}} \right) - \left( A_{ij}^{-} - \gamma \frac{s_i^{-} s_j^{-}}{v^{-}} \right) \right] 1(z_i = z_j),$$

where $1(\cdot)$ is indicator function and $\gamma > 0$ is the resolution parameter. Under this formulation, positive correlations favor assignment of ROIs to the same community, whereas negative correlations favor their separation into different communities (Rubinov & Sporns, 2010) The resolution parameter $\gamma$ controls the granularity of the detected communities, with larger values generally producing a larger number of smaller communities and smaller values producing fewer, larger communities. The Louvain Clustering algorithm is implemented through the following iterative procedure:

- Step 1: Initialize the algorithm by assigning each ROI to an individual community.
- Step 2: Sequentially consider each ROI and evaluate changes in the signed modularity objective resulting from moving the ROI to another community. The ROI is reassigned whenever the move produces a positive improvement in modularity.
- Step 3: Repeat the reassignment procedure until no additional single-node movement increases the modularity objective.
- Step 4: Aggregate the communities obtained in Step 3 into a reduced network, where each detected community is represented as a single node and the edge weights between the new nodes are obtained by combining the corresponding edge weights in the previous network.
- Step 5: Repeat Steps 2–4 on the reduced network until no additional improvement in the modularity objective can be obtained.
- Step 6: Return the resulting community assignments. The number of communities is therefore determined by the modularity optimization and the chosen value of the resolution parameter rather than being fixed in advance.

We implemented Louvain community detection using the MATLAB function '`community_louvain`' from the Brain Connectivity Toolbox (Rubinov & Sporns, 2010). We used the '`negative_sym`' objective function, which symmetrically incorporates positive and negative network weights. The resolution parameter was varied over $\gamma \in \{0,0.05,0.10,\ldots,4\}$. Because Louvain optimization is stochastic and may produce different locally optimal partitions across runs, the algorithm was repeated 1000 times for each value of $\gamma$. The resulting partitions were subsequently grouped according to the number of detected communities.

### Clustering algorithms for choosing the optimal number of communities

We use the following notations to define the metrices. We denote $A$ as the weighted adjacency matrix of the network with $n$ ROIs. Let, $d_{ij}$ be the distance/dissimilarity between ROIs $i$ and $j$ for $1 \leq i,j \leq n$ is known. We also define $C = \{C_1, \ldots C_K\}$ as the $K$ communities and $n_k$ be the size of $k^{th}$ community. We denote $z_i$ as the community label for any ROI $1 \leq i \leq n$.

#### Loglikelihood approach for WSBM

The WSBM implementation assumes that the number of communities $K$ is known apriori. However, in many real-world applications, the value of $K$ is unknown. Let $M$ denote the candidate models with $K$ communities. Then, using equation (S1), the marginal likelihood $P(A|M)$, present in the Bayes factor, is obtained by summing over the latent community assignments and integrating over the unknown community specific parameter and can be written as follows

$$P(A|M) = P(A|K) = \sum_{z} \int P(A|z,\mu,\Sigma,K)\,\pi(z,\mu,\Sigma|K)\ d\mu\ d\Sigma.$$

Following (Aicher et al., 2013, 2015), we use the variational Bayes procedure to obtain a lower bound $G_K$ on the marginal log-likelihood, $G_K \leq \log P(A|K)$. The optimized variational lower bound is then used to approximate the marginal log-likelihood. Let $M_K$ be the model containing $K$ communities and $M_{K+1}$ be the model with $K+1$ communities. We define the Bayes factor as,

$$\log B(M_K, M_{K+1}) = \log \frac{P(A|K)}{P(A|K+1)} \approx G_K - G_{K+1}.$$

Increasing $K$ enlarges both the latent assignment space and the number of block-specific parameters. Unlike the maximized conditional likelihood, the marginal likelihood averages over these unknown quantities under their priors and therefore provides an implicit, prior-dependent penalty for model complexity. In our implementation, this trade-off is approximated through the optimized variational lower bound. $G_K$.

We constructed bootstrap confidence intervals (CIs) for the Bayes factor for multiple choices of $K$. The transition at which 95% bootstrap confidence intervals of the Bayes factor overlap with the null value 0 is considered as the optimal choice for $K$ (Aicher et al., 2013, 2015; Faskowitz & Sporns, 2020). To this end, we implement the following algorithm to detect the optimal number of communities:

- Step 1: Fit WSBM to the data 1000 times for a particular value of $K$ (Fig. 2.b)
- Step 2: Repeat step 1 for varied choices of $K$.
- Step 3: For each valid[1] community assignments, bootstrap the difference between log-likelihood values for $K$ and $(K+1)$ 2000 times.
- Step 4: The minimum value of $K$ for which the 95% confidence interval for the differences overlapped with 0 is taken as the optimal number of communities (Fig. 2.c).

For each dataset, we implemented the above-described algorithm for $K = 2$ to 20 to identify the optimal number of communities. After detecting the optimal number of communities using this procedure, the final solution is chosen to be the one with maximum-log-likelihood (ML) among all valid solutions for the optimal $K$. Since the exact expression of the maximum log-likelihood is intractable, this procedure provides a close approximation to the maximum likelihood estimate (MLE) of the community assignment.

#### Silhouette Index

To implement both Spectral and K-means Clustering algorithms, a prior information on the number of clusters is needed. For both K-means Clustering and Spectral Clustering, we applied the algorithms for varied choices of $K = 2$ to 20. At each level of $K$, silhouette index (described later)

[1] Since the number of unique communities may differ from the input $K$ and due to the algorithm's time complexity (Faskowitz & Sporns, 2020), we only keep the valid solutions (out of 1000 replications) where the number of communities is exactly $K$.

and other post hoc metrics (described in the Supplement) were used to identify the optimal number of communities based on the quality of the resulting community assignments.

Silhouette index is one most commonly used cluster validation metric (Kaufman & Rousseeuw, 2009). Let $C = \{C_1, \ldots C_K\}$ be the $K$ communities and $n_k$ be the size of $k^{th}$ community. We define the average distance for each ROI $i \in C_k$ to every other ROI in the same community as, $a_i = \frac{1}{n_k - 1} \sum_{j \in \{C_k \setminus i\}} d_{ij}$ and the minimum average distance to other communities as, $b_i = \min_{s \neq k} \frac{1}{n_s} \sum_{j \in C_s} d_{ij}$. Thus, the silhouette index is defined as

$$S(k) = \frac{1}{n} \sum_{1 \le i \le n} \frac{b_i - a_i}{\max\{a_i, b_i\}}.$$

The value of silhouette index lies between $[-1,1]$. A high value indicates that the objects are well matched to their own clusters and poorly matched with members from other clusters.

#### Modularity

Modularity (Newman, 2004) measures how cohesively the ROIs of a network are grouped together, and is defined as

$$Q = \frac{1}{2m} \sum_{1 \le i,j \le n} \left( A_{ij} - \frac{d_i d_j}{2m} \right) 1(z_i = z_j),$$

where $m = \frac{1}{2} \sum_{1 \le i,j \le n} A_{ij}$ is the average edge-weights, $d_i = \sum_{1 \le j \le n; j \neq i} A_{ij}$ denotes the degree of the $i^{th}$ ROI, and $1(z_i = z_j)$ is the indicator function taking value 1 if $z_i = z_j$; 0 otherwise. A high value of modularity indicates that the network is assortative.

#### Variational Information distance

The Variational Information (VI) distance (Meilă, 2007) measures the quality of two community assignments $C^{(1)}$ and $C^{(2)}$ based on entropy. For any community assignment $C$, we define the entropy of the assignment as

$$H(C) = - \sum_{1 \le k \le K} P(k) \log P(k),$$

where $P(k) = \frac{n_k}{n}$ is the proportion of ROIs being assigned to $k^{th}$ community for $1 \le k \le K$. Now, the mutual information for two community assignments $C^{(1)}$ and $C^{(2)}$ is defined in terms of Kullback-Leibler divergence as follows,

$$I\left(C^{(1)}, C^{(2)}\right) = \sum_{1 \le k_1 \le K} \sum_{1 \le k_2 \le K} P(k_1, k_2) \log \frac{P(k_1, k_2)}{P(k_1) P(k_2)},$$

where $P(k_1, k_2) = \frac{|C_{k_1}^{(1)} \cap C_{k_2}^{(2)}|}{n}$ calculates the proportion of ROIs which are common in two different community assignments. Thus, the VI distance is defined as

$$VI\left(C^{(1)}, C^{(2)}\right) = H\left(C^{(1)}\right) + H\left(C^{(2)}\right) - I\left(C^{(1)}, C^{(2)}\right).$$

A low value of VI distance indicates that the community assignments convey similar information.

#### Calinski-Harabasz index

The Clainski-Harabasz (CH) index (Calinski & Harabasz, 1974) measures strength of dissimilarity within-group and among-group sum of squares. For a community assignment $C$, it is defined as pseudo-F ratio,

$$CH(K) = \frac{SSA/_{(K-1)}}{SSW/_{(n-K)}},$$

where the within-group sum of squares is defined as

$$SSW = \sum_{1 \le k \le K} \frac{1}{n_k} \sum_{i,j \in C_k, i<j} d_{ij}^2,$$

and the among-group sum of squares is defined as $SSA = SST - SSW$ with

$$SST = \frac{1}{n} \sum_{1 \le i < j \le n} d_{ij}^2.$$

It is evident that the value of the index will always be positive. If the ratio is very large that

indicates high separateness and compactness (Milligan & Cooper, 1985).

#### C-Index

The C-index (Milligan & Cooper, 1985) measures the within-group distance and computed as,

$$C(k) = \frac{SW - S_{min}}{S_{max} - S_{min}},$$

where the sum of within-community distances

$$SW = \sum_{1 \le k \le K} \sum_{i,j \in C_k, i<j} d_{ij}.$$

$S_{min}$ and $S_{max}$ is the sum of the $N_W$ smallest and largest distances respectively, where $N_W = \sum_{1 \le k \le K} \binom{n_k}{2}$ is the total number of ROIs is the same community. C-index lies between [0,1] and small value suggests optimal number of communities.

#### Dunn index

The Dunn index (Dunn, 1973) also measures the within-group distance and is defined as,

$$D(K) = \frac{\min_{1 \le r < s \le K} \delta(C_r, C_s)}{\max_{1 \le k \le K} \Delta_k},$$

where $\delta(C_r, C_s)$ is the inter-group distance between communities $C_r$ and $C_s$, is defined as $\delta(C_r, C_s) = \min_{i \in C_r, j \in C_s} d_{ij}$. In the denominator, the diameter of community $C_k$ is defined as $\Delta_K = \max_{i,j \in C_k} d_{ij}$. The index is maximized when clusters are compact in other words the inter-group distance is large, and the diameter is small.

#### zRand index

The zRand index (Traud et al., 2011) is a standardized pair-counting measure used to quantify the similarity between two partitions. Consider two partitions of $n$ ROIs and let $n_{ab}$ denote the number of ROIs that belong to community $a$ in the first partition and community $b$ in the second partition. Define $M = \binom{n}{2}$, as the total number of pairs of ROIs and $M_1 = \sum_a \binom{n_{a\cdot}}{2}, M_2 = \sum_b \binom{n_{\cdot b}}{2}$, where $n_{a\cdot} = \sum_b n_{ab}$ and $n_{\cdot b} = \sum_a n_{ab}$. Thus, $M_1$and $M_2$ are the numbers of ROI pairs assigned to the same community in the first and second partitions, respectively. Let $w = \sum_{a,b} \binom{n_{ab}}{2}$ denote the number of pairs assigned to the same community in both partitions. Under the null model of random assignments conditional on fixed community sizes the mean of $w$ is $\mu_w = E(w) = \frac{M_1 M_2}{M}$. Then, the zRand index is defined as

$$z_R = \frac{w - \mu_w}{\sigma_w},$$

where $\sigma_w^2 = Var(w)$. Following (Traud et al., 2011), the analytical variance is

$$\sigma_w^2 = \frac{M}{16} - \frac{(4M_1 - 2M)^2 (4M_2 - 2M)^2}{256M^2} + \frac{C_1 C_2}{16n(n-1)(n-2)} + \frac{[(4M_1 - 2M)^2 - 4C_1 - 4M][(4M_2 - 2M)^2 - 4C_2 - 4M]}{64n(n-1)(n-2)(n-3)},$$

where $C_1 = n(n^2 - 3n - 2) - 8(n+1)M_1 + 4\sum_a n_{a\cdot}^3$ and $C_2 = n(n^2 - 3n - 2) - 8(n+1)M_2 + 4\sum_b n_{\cdot b}^3$. A large positive value of $z_R$ indicates that the two partitions have substantially more pairs of ROIs assigned together than would be expected under the random-assignment null model. A value near zero indicates similarity close to the null expectation, whereas a negative value indicates less agreement than expected by chance. Importantly, $z_R$ is a standardized similarity measure and is not restricted to the interval [0,1].

### Network topology

Consider a network consisting of $K$ communities or block where $(\mu_{ab})$ denotes the $K \times K$ mean connectivity matrix among the networks. Similar to (Faskowitz & Sporns, 2020), communities can be classified into different categories depending on the connectivity patterns as described below.

(i) On-diagonal: blocks representing within-community connectivity, i.e, $\mu_{ab}$ when $a = b$.

(ii) Assortative: minimum on-diagonal value is larger than the off-diagonal value, i.e, $\min(\mu_{aa}, \mu_{bb}) > \mu_{ab}$.

(iii) Core/periphery: off-diagonal value is larger than one on-diagonal value; if the off-diagonal value is closer to the larger of the on-diagonal values, it is core; otherwise it is periphery, i.e., $\mu_{aa} > \mu_{ab} > \mu_{bb}$ or $\mu_{bb} > \mu_{ab} > \mu_{aa}$.

(iv) Disassortative: off-diagonal value is larger than on-diagonal values, i.e., $\mu_{ab} > \max(\mu_{aa}, \mu_{bb})$.

(v) Unspecified: do not belong to any of the above described topology structures.

Tables S3-S5 report the proportion of all topological structures across different atlases for the HCP, WU120 and BCP datasets respectively. For any atlas with $K$ communities, the proportions are computed from average connectivity pattern of the FC matrix over all possible $\frac{K(K+1)}{2}$ community-community interactions.

**Table S1:** Simulation setting

| $K$ | **Community sizes** | **ROI** |
|---|---|---|
| 5 | 30, 15, 25, 20, 10 | 100 |
| 7 | 40, 25, 30, 20, 35, 30, 20 | 200 |
| 10 | 30, 18, 20, 15, 17, 22, 18, 21, 17, 22 | 200 |

**Table S2:** Runtime complexity of algorithms

| **Algorithm** | **Time complexity** |
|---|---|
| WSBM | $O(n^2K^2)$ |
| Spectral Clustering | $O(n+K)$ |
| K-means Clustering | $O(n+K)$ |
| Consensus Algorithm | $O(n+K^3)$ |

**Table S3**: Proportion of different network topologies for HCP dataset across different atlases.

| | **WSBM (MLE)** | **Spectral Clustering** | **K-means Clustering** | **WSBM (Consensus)** | | | |
|---|---|---|---|---|---|---|---|
| ***K*** | 11 | 11 | 11 | 11 | 12 | 13 | 14 |
| On-diagonal | 0.167 | 0.167 | 0.167 | 0.167 | 0.154 | 0.143 | 0.133 |
| Assortative | 0.803 | 0.833 | 0.833 | 0.833 | 0.833 | 0.835 | 0.724 |
| Core | 0 | 0 | 0 | 0 | 0 | 0 | 0 |
| Periphery | 0.030 | 0 | 0 | 0 | 0.013 | 0.022 | 0.019 |
| Disassortative | 0 | 0 | 0 | 0 | 0 | 0 | 0 |
| Unspecified | 0 | 0 | 0 | 0 | 0 | 0 | 0.124 |

**Table S4**: Proportion of different network topologies for WU 120 dataset across different atlases.

| | **WSBM (MLE)** | **Spectral Clustering** | **K-means Clustering** | **WSBM (Consensus)** | | |
|---|---|---|---|---|---|---|
| ***K*** | 11 | 11 | 11 | 11 | 13 | 14 |
| On-diagonal | 0.167 | 0.167 | 0.167 | 0.167 | 0.143 | 0.133 |
| Assortative | 0.833 | 0.833 | 0.833 | 0.833 | 0.857 | 0.867 |
| Core | 0 | 0 | 0 | 0 | 0 | 0 |
| Periphery | 0 | 0 | 0 | 0 | 0 | 0 |
| Disassortative | 0 | 0 | 0 | 0 | 0 | 0 |
| Unspecified | 0 | 0 | 0 | 0 | 0 | 0 |

**Table S5**: Proportion of different network topologies for BCP dataset across different atlases.

| | **WSBM (MLE)** | **Spectral Clustering** | **K-means Clustering** | **WSBM (Consensus)** | | | |
|---|---|---|---|---|---|---|---|
| ***K*** | 15 | 15 | 15 | 13 | 14 | 15 | 16 |
| On-diagonal | 0.125 | 0.125 | 0.125 | 0.143 | 0.133 | 0.125 | 0.118 |
| Assortative | 0.875 | 0.875 | 0.875 | 0.857 | 0.867 | 0.875 | 0.882 |
| Core | 0 | 0 | 0 | 0 | 0 | 0 | 0 |
| Periphery | 0 | 0 | 0 | 0 | 0 | 0 | 0 |
| Disassortative | 0 | 0 | 0 | 0 | 0 | 0 | 0 |
| Unspecified | 0 | 0 | 0 | 0 | 0 | 0 | 0 |

**Table S6:** The number of valid solutions recovered for WSBM in adult and infant datasets.

| ***K*** | **2-4** | **5** | **6** | **7** | **8** | **9** | **10** | **11** | **12** | **13** | **14** | **15** | **16** | **17** | **18** |
|---|---|---|---|---|---|---|---|---|---|---|---|---|---|---|---|
| **HCP** | 1000 | 996 | 984 | 903 | 787 | 585 | 383 | 201 | 107 | 48 | 18 | 6 | 3 | 0 | 0 |
| **WU 120** | 1000 | 998 | 976 | 912 | 744 | 557 | 360 | 177 | 102 | 50 | 14 | 6 | 1 | 0 | 0 |
| **BCP** | 1000 | 1000 | 1000 | 995 | 958 | 868 | 736 | 546 | 392 | 237 | 118 | 57 | 27 | 8 | 1 |

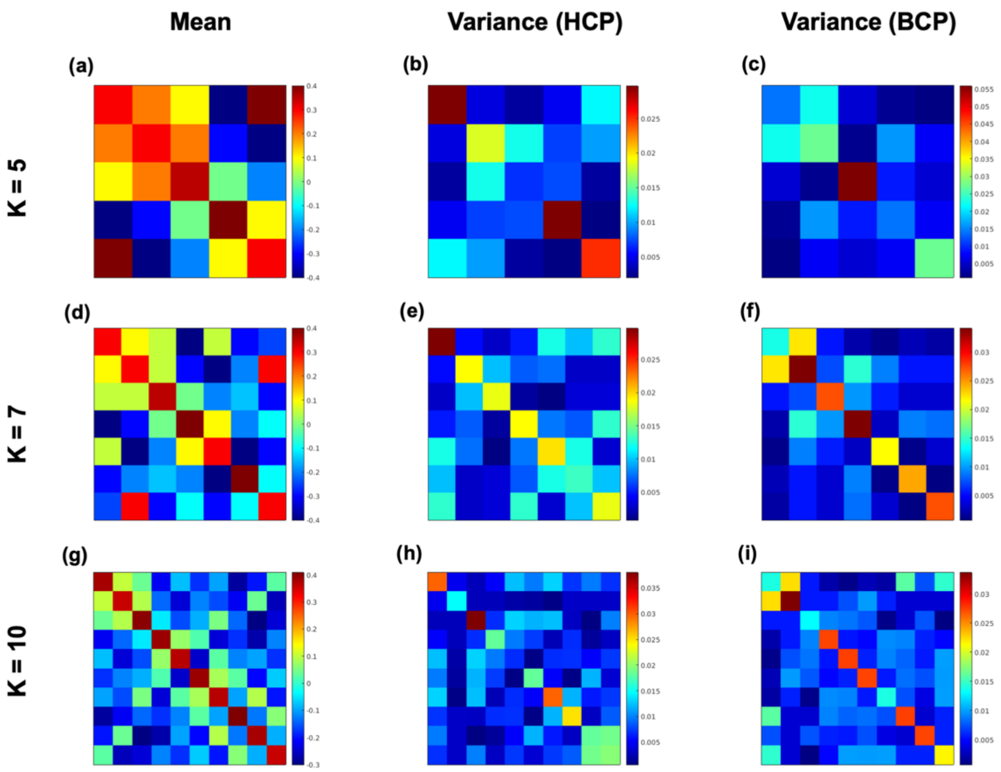


**Figure S1.** Synthetic network means and variances. **(a)** Average community connectivity matrix. **(b)** Variability across the community.

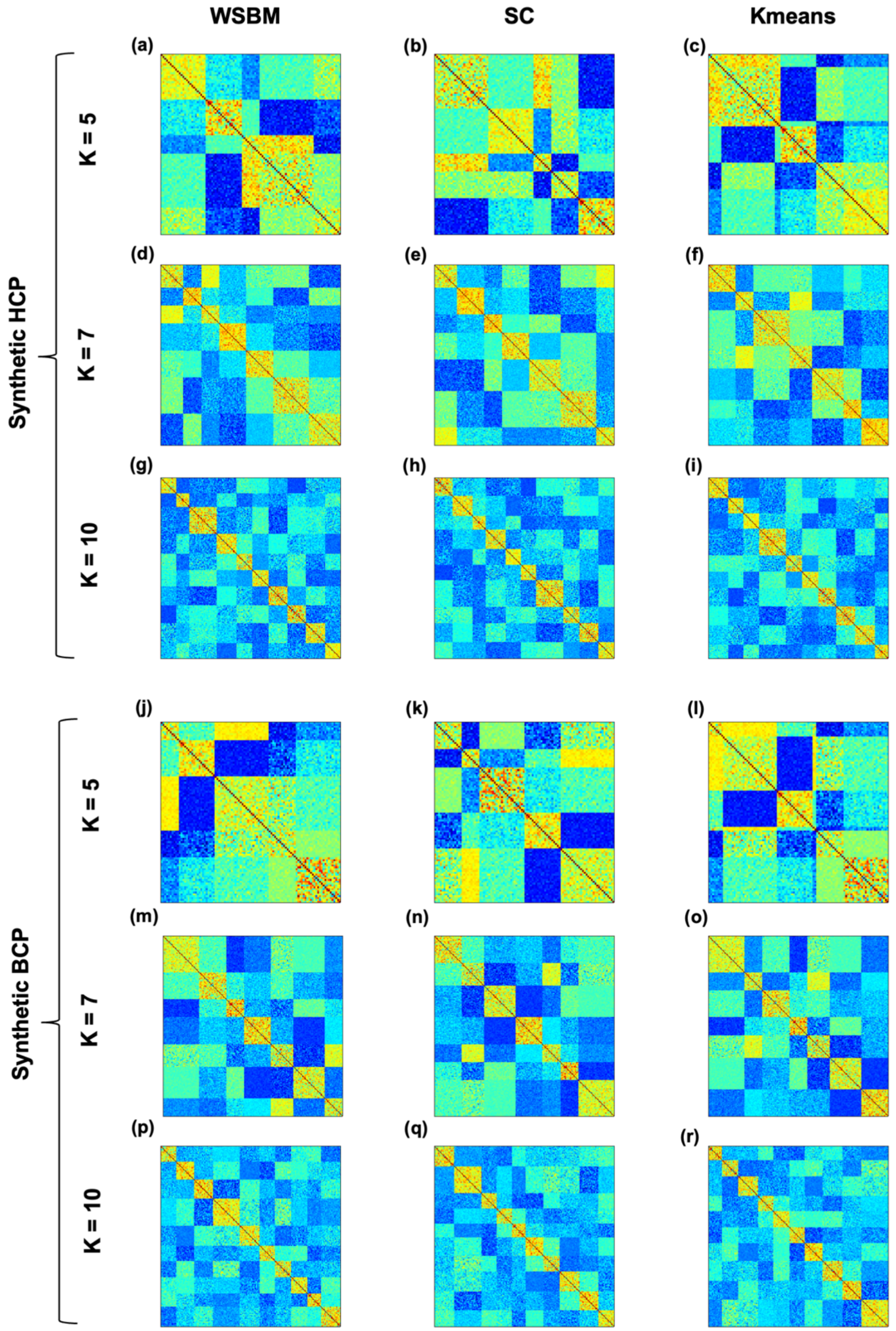


**Figure S2.** Illustration of community detection solutions for one randomly selected synthetic functional connectivity (FC) matrix out of 100 independently simulated matrices for each setting, as presented in Fig. 1. Community-ordered FC matrices are shown for WSBM, Spectral Clustering (SC), and K-means clustering at $K = 5{,}7$ and 10 communities. The top three rows **(a–i)** correspond to HCP and the bottom three rows **(j–r)** to BCP. Within each matrix, ROIs are reordered according to their estimated community assignments, such that contiguous diagonal blocks represent ROIs assigned to the same community. Columns correspond to WSBM, SC, and K-means, respectively.

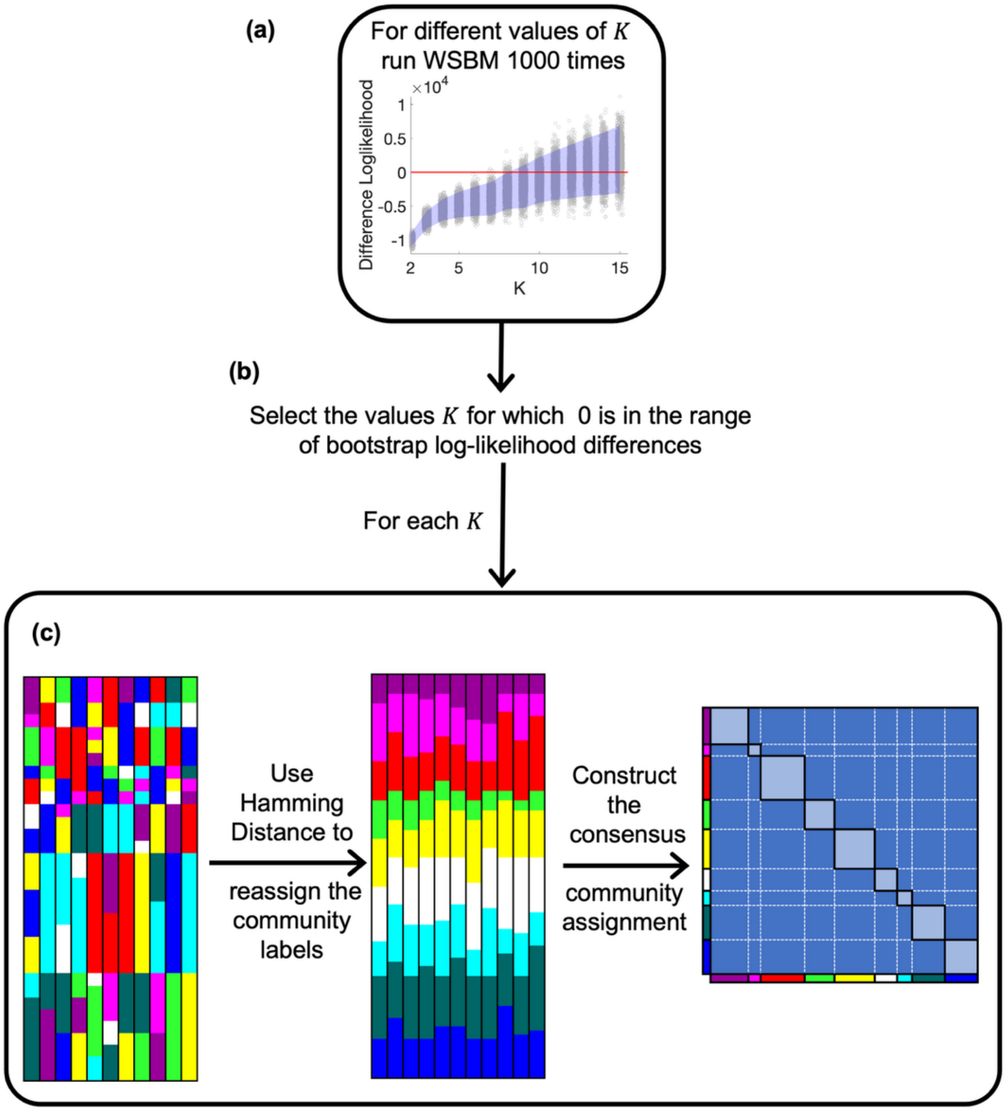


**Figure S3.** WSBM consensus pipeline. **(a)** Bootstrap difference log-likelihood plot for WSBM. **(b)** Choice of $K$ for consensus. (c) Consensus community assignment for a given $K$.

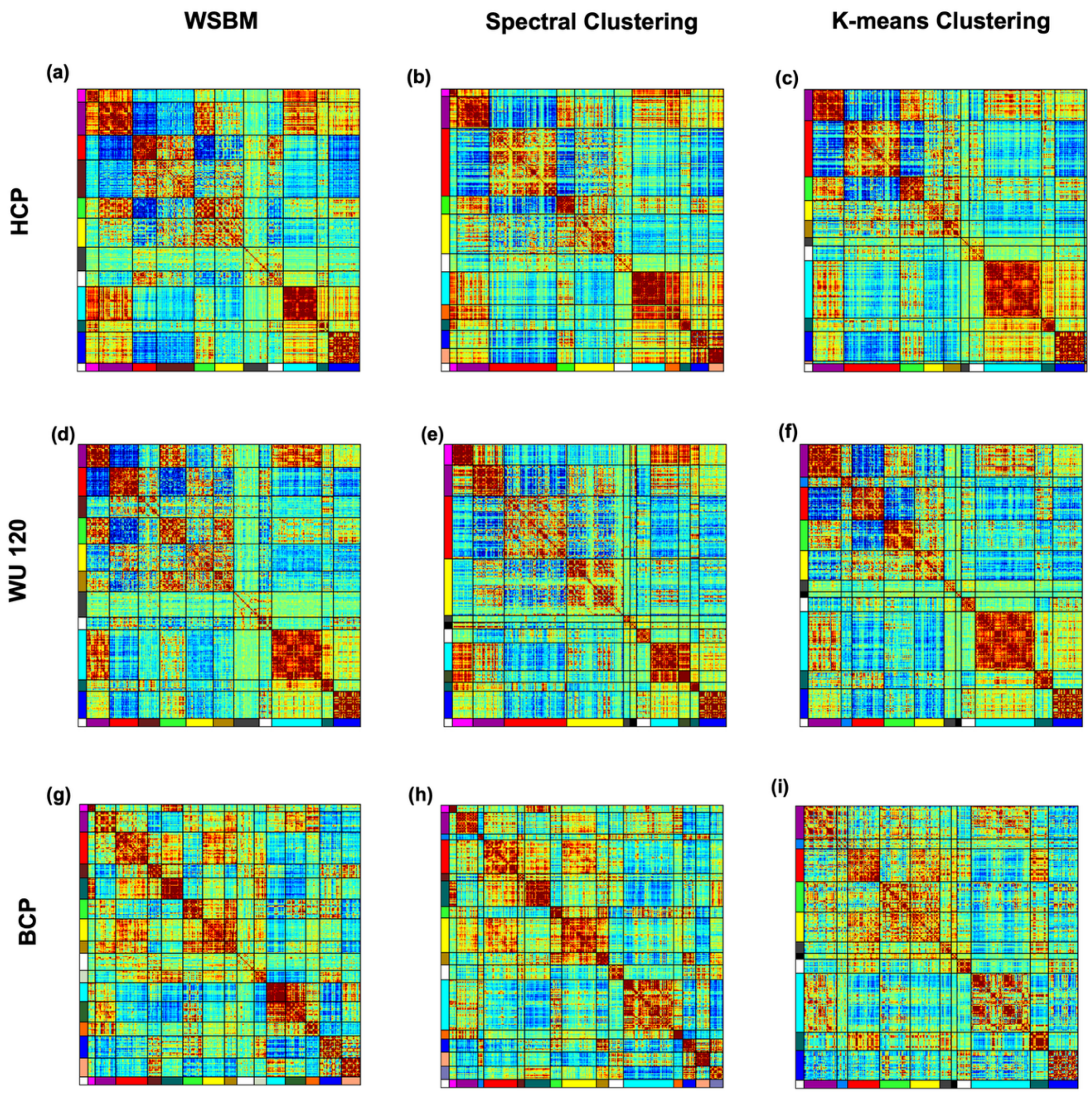


**Figure S4.** Average FC matrices plots sorted according to the community labels. **(a-f)** Average FC matrix plots for adult datasets with 11 communities. **(g-i)** Average FC matrix plots for infant dataset with 15 communities.

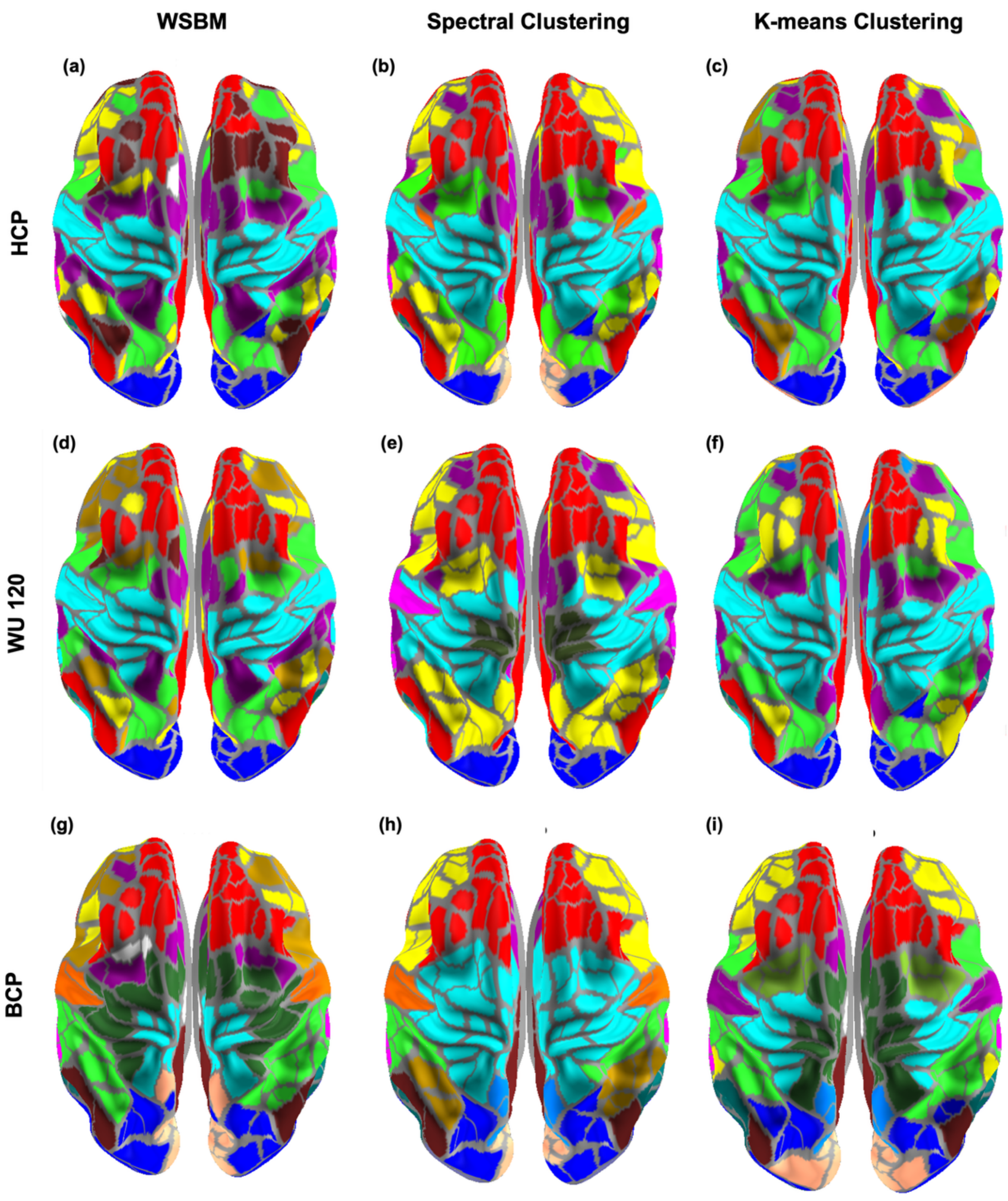


**Figure S5.** Axial views of the brain surface plots. **(a-f)** Brain Surface plots for adult datasets with 11 communities. **(g-i)** Brain surface plots infant dataset with 15 communities.

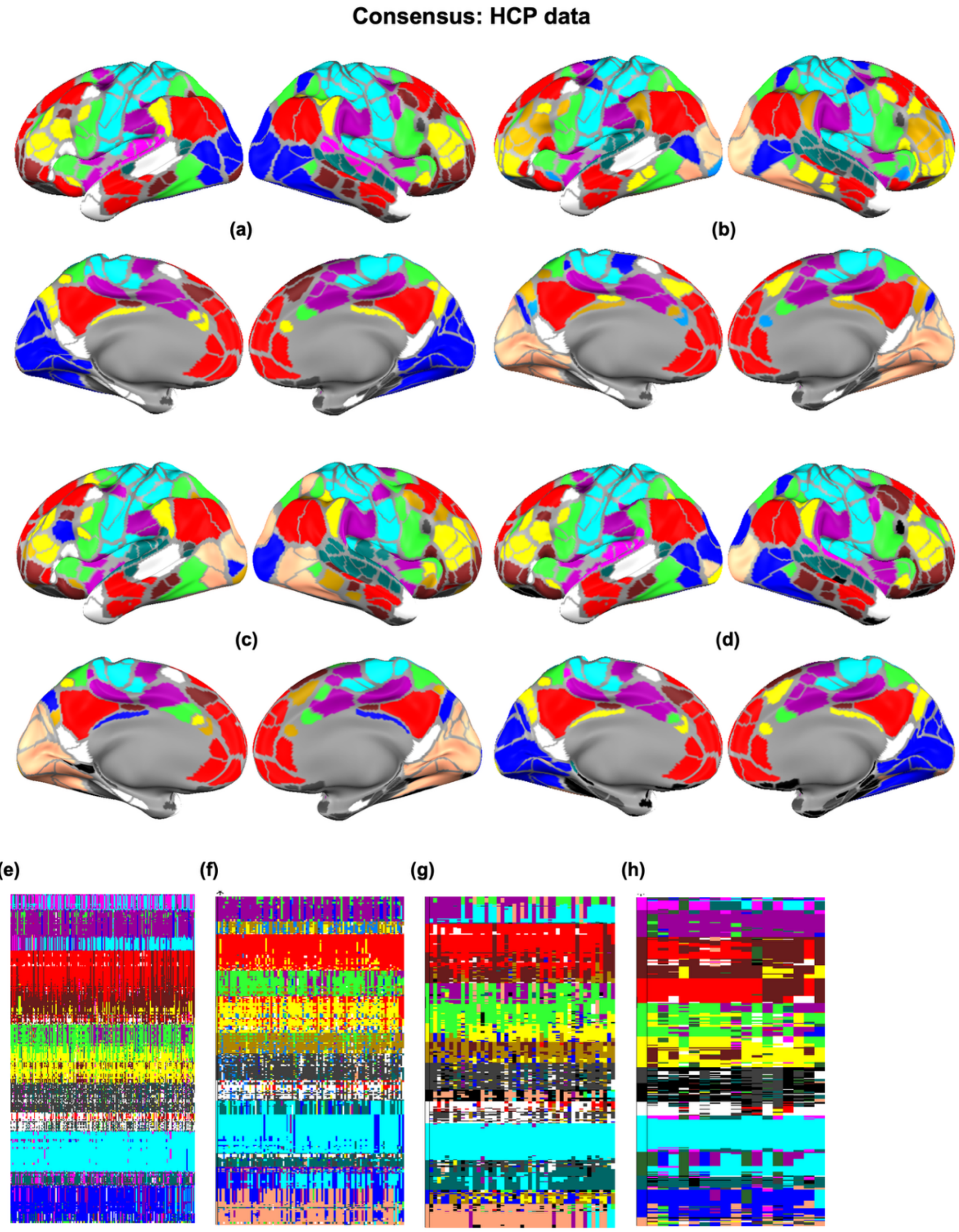


**Figure S6.** Consensus solutions for HCP dataset. **(a-d)** Brain surface plots of consensus for WSBM on HCP data with $K$ = 11,12, 13 and 14 respectively. **(e-h)** Community assignments for valid replications of WSBM on HCP data with $K$ = 11,12, 13 and 14 respectively.

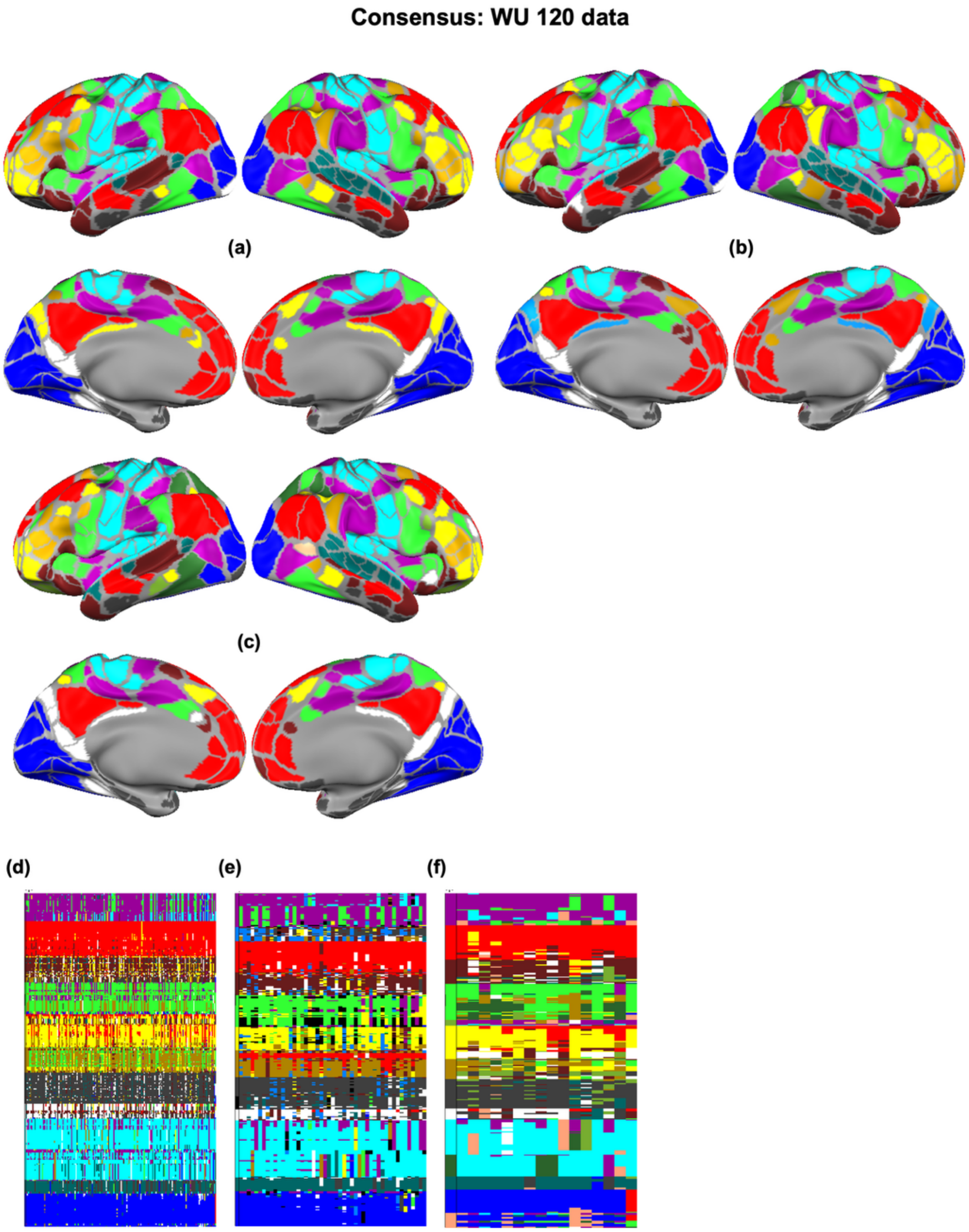


**Figure S7.** Consensus solutions for WU 120 dataset. **(a-c)** Brain surface plots of consensus for WSBM on WashU 120 data with $K$ = 11,13 and 14 respectively. **(d-f)** Community assignments for valid replications of WSBM on WU 120 data with $K$ = 11,13 and 14 respectively.

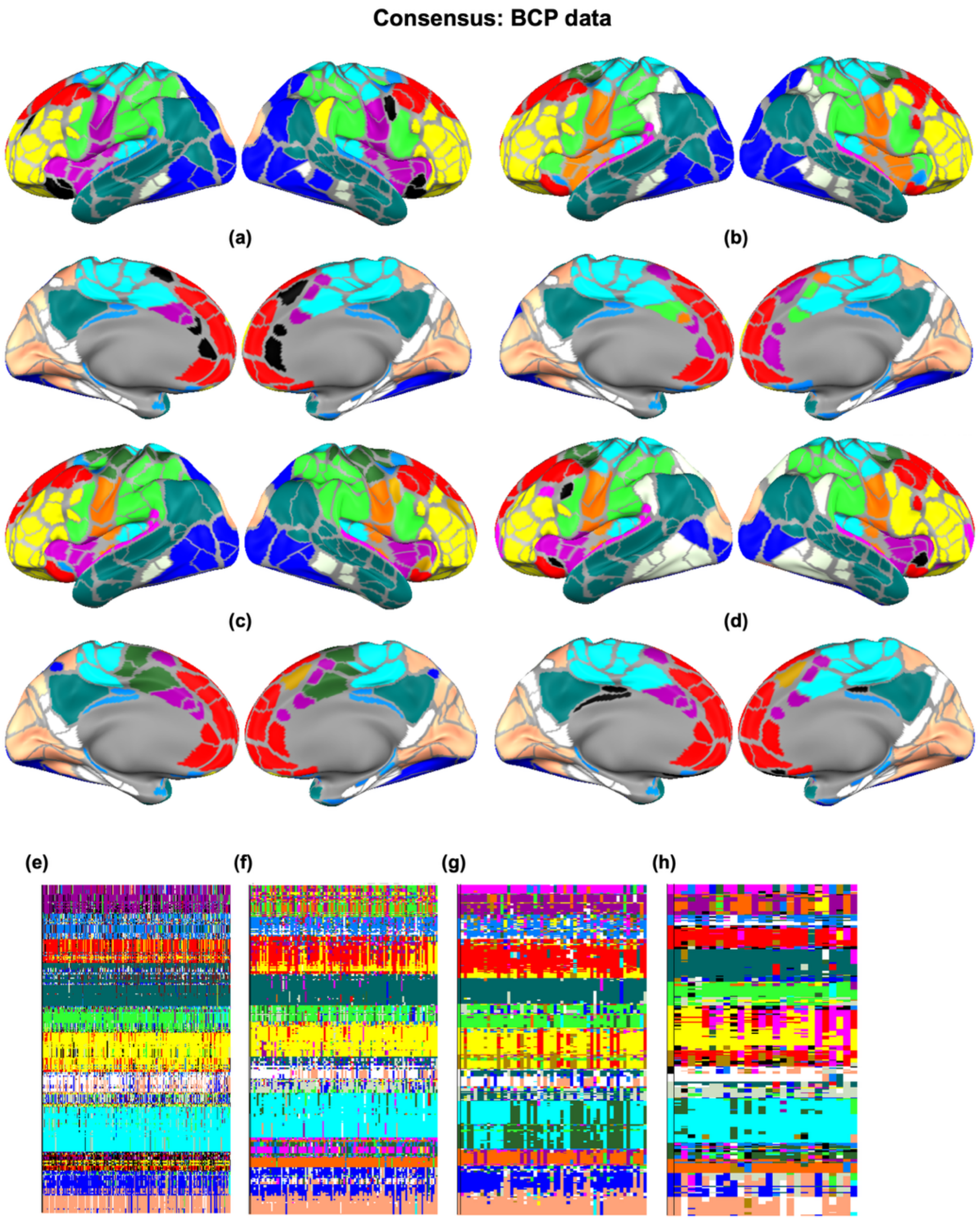


**Figure S8.** Consensus solutions for BCP dataset. **(a-d)** Brain surface plots of consensus for WSBM on BCP data with $K = 13,14,15$ and 16 respectively. **(e-f)** Community assignments for valid replications of WSBM on BCP data with $K = 13,14,15$ and 16 respectively.

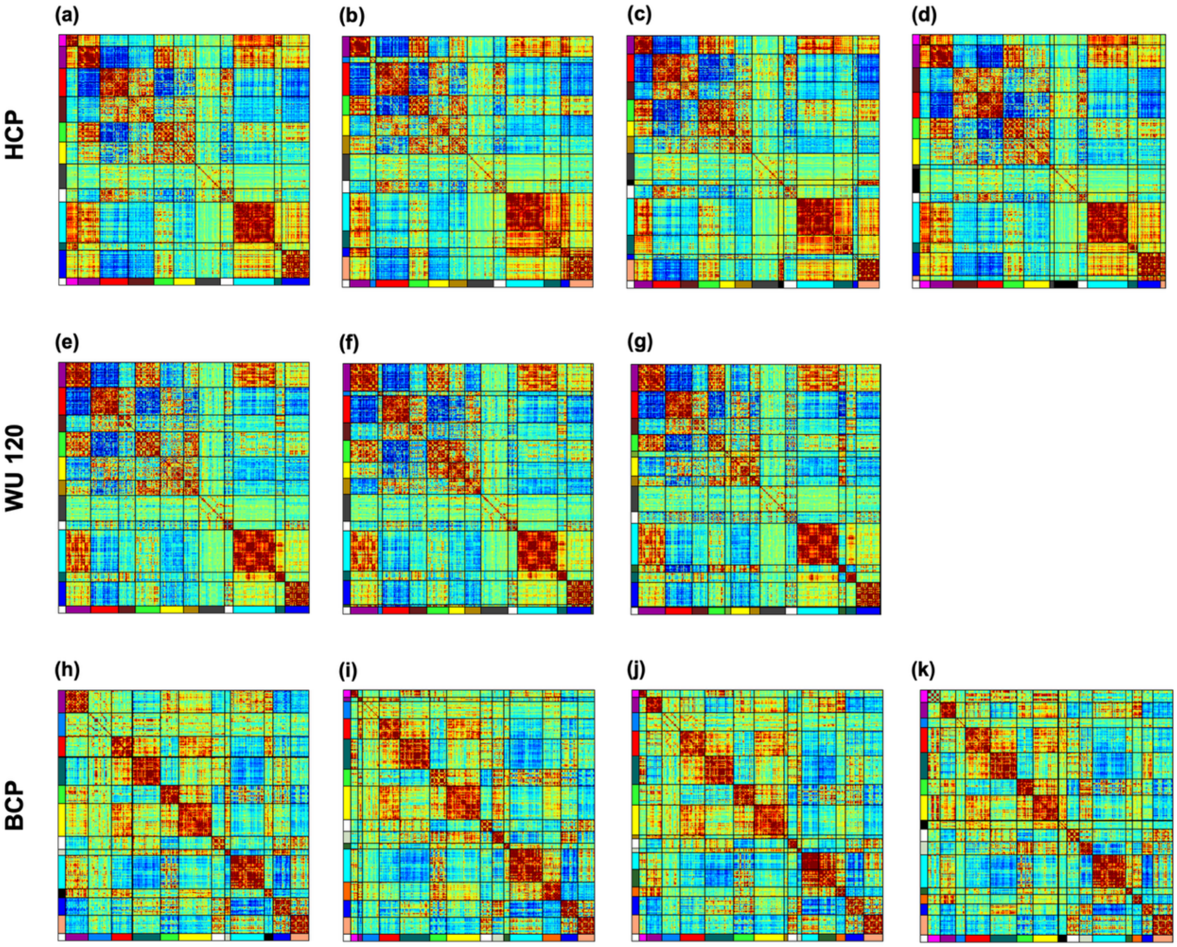


**Figure S9.** Average FC matrix plots for the consensus solution of adults and infant datasets. **(a-d)** Average FC matrix plots of consensus for WSBM on HCP data with $K$ = 11, 12, 13 and 14 respectively. **(e-g)** Average FC matrix plots of consensus for WSBM on WU 120 data with $K$ = 11, 13 and 14 respectively. **(h-k)** Average FC matrix plots of consensus for WSBM on BCP data with $K$ = 13, 14, 15 and 16 respectively.

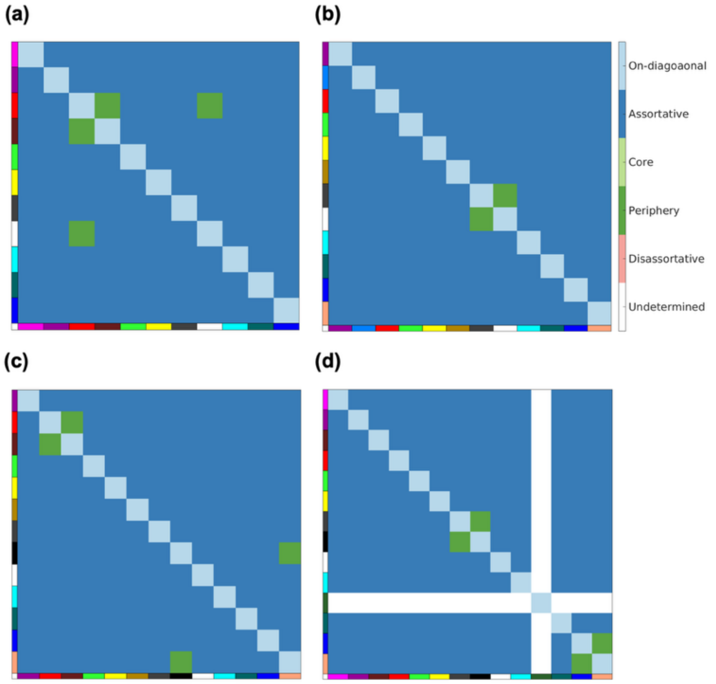


**Figure S10.** Network topology for HCP data. **(a)** WSBM (MLE) solution for $K$=11. **(b)** WSBM (Consensus) solution for $K$=12. **(c)** WSBM (Consensus) solution for $K$=13, **(d)** WSBM (Consensus) solution for $K$=14.
